# TCR signaling promotes the assembly of RanBP2/RanGAP1-SUMO1/Ubc9 nuclear pore subcomplex via PKC-θ-mediated phosphorylation of RanGAP1

**DOI:** 10.1101/2021.03.10.434793

**Authors:** Yujiao He, Zhiguo Yang, Chen-Si Zhao, Yu Gong, Zhihui Xiao, Yun-Yi Li, Yiqi Chen, Dianying Feng, Amnon Altman, Yingqiu Li

**Author notes:** Correspondence should be addressed to Y.L.

## Abstract

The nuclear pore complex (NPC) is the sole and selective gateway for nuclear transport and its dysfunction has been associated with many diseases. The NPC subcomplex RanBP2, which consists of RanBP2 (Nup358), RanGAP1-SUMO1 and Ubc9, regulates the assembly and function of the NPC. The roles of immune signaling in NPC assembly remain poorly understood. Here, we show that, following TCR stimulation, protein kinase C-θ (PKC-θ) directly phosphorylates RanGAP1 to facilitate RanBP2 subcomplex assembly and nuclear import and, thus, the nuclear translocation of AP-1 transcription factor. Mechanistically, TCR stimulation induces the translocation of activated PKC-θ to the NPC, where it interacts with and phosphorylates RanGAP1 on Ser^504^ and Ser^506^. RanGAP1 phosphorylation increases its binding affinity for Ubc9, thereby promoting sumoylation of RanGAP1 and, finally, assembly of the RanBP2 subcomplex. Our findings reveal an unexpected role of PKC-θ as a direct regulator of nuclear import and uncover a phosphorylation-dependent sumoylation of RanGAP1, delineating a novel link between TCR signaling and assembly of the RanBP2 NPC subcomplex.

## Introduction

Nuclear pore complexes (NPCs) span the nuclear envelope (NE) and mediate nucleo-cytoplasmic exchange. NPC dysfunction has been associated with various human diseases(Beck & Hurt, 2017). NPCs in the animal kingdom are ~110 megadalton supramolecular assemblies of multiple copies of ~30 different nuclear pore proteins, termed nucleoporins (NUPs)(Hampoelz, Andres-Pons, Kastritis, & Beck, 2019; D. H. Lin & Hoelz, 2019). The elaborate structure of NPCs consists of several biochemically and ultrastructurally defined substructures, namely, cytoplasmic filaments, cytoplasmic and nuclear rings, the inner pore ring, the central transporter region, and the nuclear basket(Hampoelz, Andres-Pons, et al., 2019; D. H. Lin & Hoelz, 2019; Otsuka & Ellenberg, 2018).

An important component of the cytoplasmic filaments is the RanBP2 subcomplex that consists of RanBP2 (Ran-binding protein 2, Nup358), RanGAP1 (Ran GTPase-activating protein 1) SUMO1 and Ubc9 (SUMO-conjugating enzyme)(Hampoelz, Andres-Pons, et al., 2019; D. H. Lin & Hoelz, 2019; Werner, Flotho, & Melchior, 2012). This subcomplex has multiple functions in nucleo-cytoplasmic transport by coordinating importin β recycling and facilitating importin β family-mediated nuclear import(Hampoelz, Andres-Pons, et al., 2019; Hutten, Flotho, Melchior, & Kehlenbach, 2008; D. H. Lin & Hoelz, 2019). serving as a disassembly machine for export complexes(Ritterhoff et al., 2016), stabilizing the interaction between the inner and outer Y complex of the cytoplasmic ring in humans(von Appen et al., 2015), and controlling NPC assembly beyond nuclei during oogenesis(Hampoelz, Schwarz, et al., 2019). RanGAP1 has two forms, *i.e.,* nonsumoylated RanGAP1 that mainly localizes in the cytoplasm, and SUMO1-conjugated RanGAP1 that resides at NPCs(Matunis, Coutavas, & Blobel, 1996), generated by Ubc9-mediated sumoylation(Bernier-Villamor, Sampson, Matunis, & Lima, 2002; Lee et al., 1998). SUMO1 conjugation of RanGAP1 is required for assembly of the RanBP2 subcomplex(Hampoelz, Andres-Pons, et al., 2019; Hampoelz, Schwarz, et al., 2019; Hutten et al., 2008; Joseph, Liu, Jablonski, Yen, & Dasso, 2004; Mahajan, Delphin, Guan, Gerace, & Melchior, 1997; Reverter & Lima, 2005; Ritterhoff et al., 2016; von Appen et al., 2015; Werner et al., 2012). However, it is unclear whether receptor signaling and, particularly, immune cell stimuli, regulate RanBP2 subcomplex assembly.

Protein kinase C-θ (PKC-θ) is a pivotal regulator of T-cell activation. PKC-θ belongs to the novel, calcium-independent subfamily of the PKC enzyme family, and is highly expressed in hematopoietic cells, particularly in T cells(Baier et al., 1993). Engagement of the TCR together with costimulatory receptors *(e.g.,* CD28) by cognate antigens and costimulatory ligands presented by antigen-presenting cells (APCs) recruits PKC-θ to the center of the T cell immunological synapse (IS) formed at the T cell-APC contact site, where it mediates activation of the transcription factors NF-κB, AP-1 and NFAT, leading to T-cell activation, cytokine (*e.g*., IL-2) production, and acquisition of effector functions(Baier-Bitterlich et al., 1996; Coudronniere, Villalba, Englund, & Altman, 2000; X. Lin, O’Mahony, Mu, Geleziunas, & Greene, 2000; Pfeifhofer et al., 2003; Wang et al., 2015; Xie et al., 2019).

AP-1 transcription factors have pleiotropic effects, including a central role in different aspects of the immune system such as T-cell activation, Th differentiation, T cell anergy and exhaustion(Atsaves, Leventaki, Rassidakis, & Claret, 2019). To date, unlike the regulation of NF-κB, which has been studied extensively, the regulation of AP-1 by PKC-θ in response to TCR stimulation is incompletely understood. Paradoxically, although PKC-θ deficiency does not impair the TCR-induced activation of c-Jun N-terminal kinase (JNK) nor the expression level of AP-1 in mature T cells, it severely inhibits the transcriptional activity of AP-1(Pfeifhofer et al., 2003; Sun et al., 2000), suggesting the existence of an unknown regulatory layer of TCR-PKC-θ signaling. Hence, further exploring how PKC-θ mediates TCR-induced AP-1 activation is important for understanding T cell immunity.

In this study, we reveal that PKC-θ directly regulates the nuclear import function of the NPC, which accounts for the effective, TCR-induced activation of AP-1. We show that PKC-θ promotes the nuclear import process by facilitating the assembly of the RanBP2 subcomplex in T cells. This nuclear import was significantly impaired in PKC-θ-deficient (*Prkcq*^-/-^ or KO) T cells. Mechanistically, PKC-θ directly interacted with and phosphorylated RanGAP1 at Ser^504^ and Ser^506^. RanGAP1 phosphorylation at these two Ser residues, but not sumoylation at Lys^524^, promotes its binding to Ubc9, thereby facilitating the sumoylation of RanGAP1 and, consequently, RanBP2 subcomplex assembly. We further demonstrate that a non-phosphorylatable double mutant (S504A and S506A; AA) of RanGAP1 displayed a significantly reduced sumoylation and blocked TCR-induced nuclear translocation of AP-1, as well as that of NF-κB and NFAT. Thus, our study demonstrates a novel PKC-θ-mediated, phosphorylation-dependent sumoylation of RanGAP1 as an obligatory step for proper assembly of the RanBP2 subcomplex and, furthermore, establishes a mechanism for the regulation of AP-1 activation by PKC-θ.

## Results

### PKC-θ translocates to the NE and colocalizes with NPCs upon TCR stimulation

Several PKC isoforms translocate to the NE after phorbol ester (PMA, a diacylglycerol mimetic and PKC activator) treatment(Collas, 1999; Leach, Powers, Ruff, Jaken, & Kaufmann, 1989). We therefore examined whether PKC-θ also translocates to the NE in stimulated Jurkat E6.1 T cells, a human leukemic T-cell line widely used for the study of TCR signaling(Abraham & Weiss, 2004). By biochemically isolating the NE, we observed that anti-CD3 plus anti-CD28 antibody (Ab) costimulation or stimulation with APCs that were pulsed with a superantigen, staphylococcal enterotoxin E (SEE), promoted PKC-θ translocation to the NE fraction (Figure 1A,B, respectively). Consistent with previous findings, TCR stimulation also promoted PKC-θ translocation to the plasma membrane (PM) fraction (Figure 1A,B). In addition, transmission electron microscopy (TEM) analysis of immunogold-labeled PKC-θ showed its translocation to the cytoplasmic face of the NE in response to anti-CD3 plus anti-CD28 costimulation (Figure 1—figure supplement 1A). Quantitation showed that after 15 min of stimulation, >20% of total PKC-θ was localized 100 nm or less from the NE, and >30% of total PKC-θ was localized 100 nm or less from the PM (Figure 1—figure supplement 1B).

**Figure 1.**
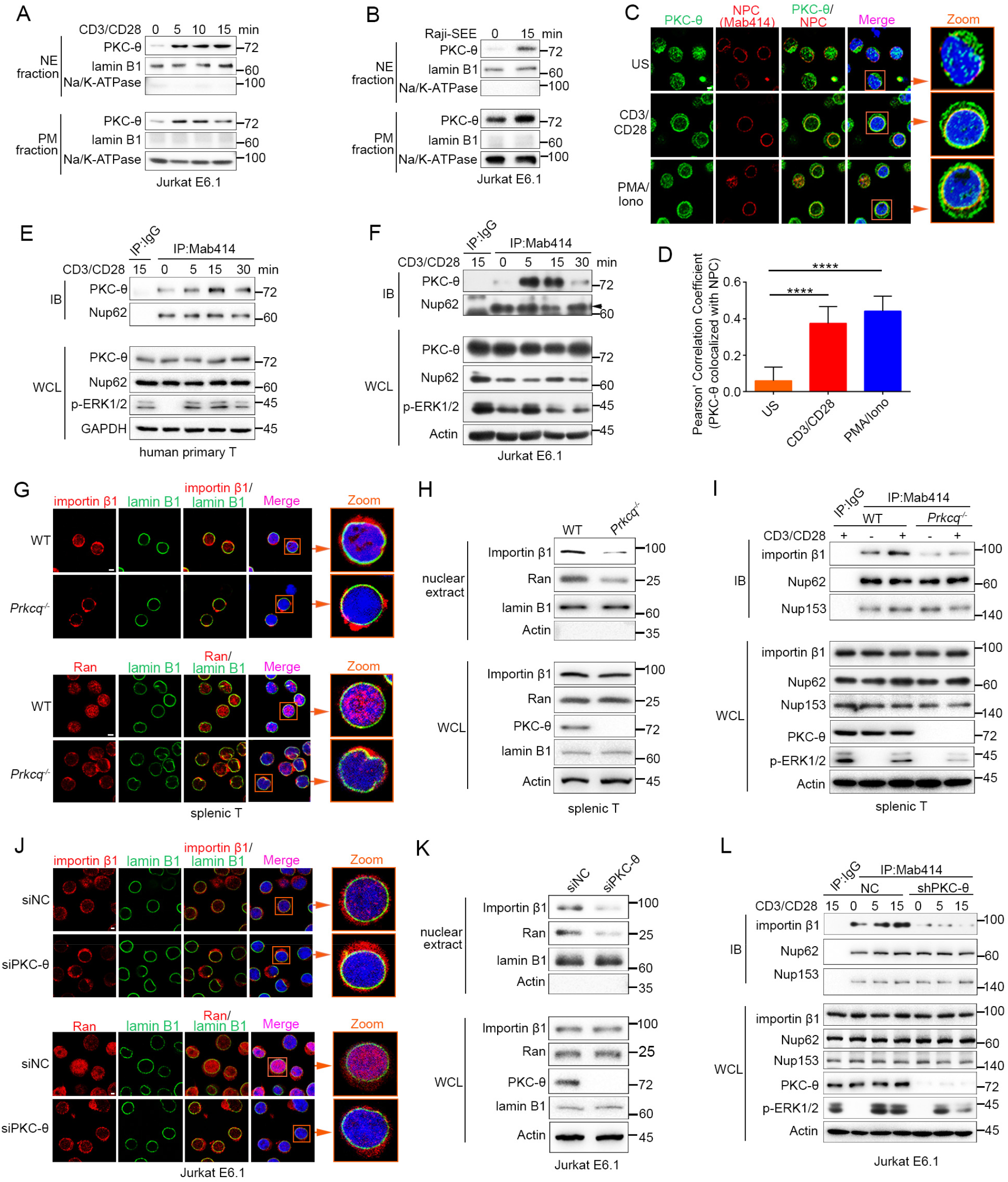
TCR stimulation promotes PKC-θ colocalization with the NPC and PKC-θ deficiency decreases nuclear import of importin β1 and Ran and NPC association with importin β1. (**A, B**) Subcellular fractionation of Jurkat E6.1 cells stimulated for 0-15 min with anti-CD3 plus anti-CD28 (A) or with superantigen (SEE)-pulsed Raji B cells (B) and immunodetection with the indicated antibodies. NE, nuclear envelope; PM, plasma membrane. (**C**) Confocal imaging of PKC-θ (green) and NPCs (Mab414, red) colocalization in representative Jurkat E6.1 cells left unstimulated (US) or stimulated for 15 min with anti-CD3 plus anti-CD28 or with PMA plus Iono. Nuclei are stained with DAPI (blue). Areas outlined by squares in the merged images are enlarged at right. Scale bars, 2 μm. (**D**) Quantification of PKC-θ colocalization with NPCs by Pearson correlation coefficient. Analysis was based on at least three different images covering dozens of cells using the ImageJ software. \*\*\*\**P* < 0.0001 (one-way ANOVA with post hoc test). (**E, F**) Immunoblot analysis of NPC IPs (Mab414) or whole cell lysates (WCL) from human primary T cells (E) or Jurkat E6.1 cells (F) stimulated for 0-30 min with anti-CD3 plus anti-CD28. Control IP with normal IgG is shown in the left lane. Nup62, an NPC component, was used a as loading control for the IPs. The arrowhead indicates the Nup62 protein band. (**G**) Confocal imaging of importin β1 and Ran in representative wild-type (WT) or *Prkcq*^-/-^ mouse primary splenic T cells stained with the indicated antibodies. Areas outlined by squares in the merged images are enlarged at right. Scale bars, 2 μm. (**H**) Subcellular fractionation of mouse splenic T cells and immunodetection with the indicated antibodies. (**I**) Immunoblot analysis of NPC IPs (Mab414) or whole cell lysates (WCL) from unstimulated or anti-CD3 plus anti-CD28-stimulated WT or *Prkcq*^-/-^ mouse splenic T cells. Control IP with normal IgG is shown in the left lane. (**J, K**) Confocal imaging of importin β1 and Ran (J) and subcellular fractionation (K), analyzed as in (G, H), of Jurkat E6.1 cells transfected with scrambled siRNA negative control (siNC) or PKC-θ targeting siRNA (siPKC-θ). Scale bars, 2 μm. (**L**) Immunoblot analysis of NPC IPs (Mab414) or whole cell lysates (WCL) from unstimulated or stimulated Jurkat E6.1 T cells stably expressing a control small hairpin RNA (shRNA) or a PKC-θ targeting shRNA (shPKC-θ), analyzed as in (I). Data are representative of three (**A, B, E, F, H, I, K, L**) or two (**C, D, G, J**) biological replicates. **Figure supplement 1.** PKC-θ translocates to the NE following TCR stimulation and PKC-θ deficiency decreases nuclear import of importin β1 and Ran and NPC association with importin β1.

Given that NPCs are the important components of the NE, we next determined whether PKC-θ colocalized with NPCs. We stimulated Jurkat T cells with anti-CD3 plus anti-CD28 Abs or with PMA plus a Ca^2+^ ionophore, ionomycin (Iono), for 15 min and imaged the cells by confocal microscopy. Staining with an anti-PKC-θ Ab and a monoclonal Ab (Mab414) that recognizes a subset of NPC proteins revealed that stimulation significantly increased PKC-θ colocalization with NPCs (Figure 1C,D). Similarly, T cells stimulated with SEE-pulsed APCs also displayed partial PKC-θ colocalization with the NPCs (Figure 1 — figure supplement 1C). Immunoprecipitation (IP) with Mab414 showed that costimulation increased PKC-θ binding to NPCs in primary human T cells (Figure 1E) and in Jurkat T cells (Figure 1F). When we stimulated T cells with PMA plus Iono, a portion of PKC-θ molecules translocated to the NE (Figure 1—figure supplement 1D-F). Together, these results demonstrate that TCR stimulation induces PKC-θ translocation to the NE and, more specifically, colocalization and physical association with NPCs, suggesting that PKC-θ may participate in the process of nucleo-cytoplasmic transport.

### PKC-θ deficiency decreases NPC association of importin β1

Importin β1 and Ran proteins play a key role in nucleo-cytoplasmic transport, and importin β1 can be specifically recruited to NPCs and then mediate the passage of proteins, including transcription factors such as AP-1, NF-κB and NFAT, into the nucleus through NPCs(Tetenbaum-Novatt & Rout, 2010; van der Watt et al., 2016). To determine the physiological relevance of the PKC-θ-NPC association, we first analyzed the cellular distribution of importin β1 and Ran in resting T cells from wild-type and *Prkcq*^-/-^ mice. Using confocal microscopy, we found that PKC-θ deletion resulted in a decreased ratio of nuclear-to-cytoplasmic of importin β1 and Ran (Figure 1G, Figure 1—figure supplement 1G). PKC-θ deletion also resulted in a decreased translocation of importin β 1 and Ran to the nuclear fraction (Figure 1H, Figure 1—figure supplement 1H). Next, we assessed the binding of importin β1 to NPCs. In wild-type T cells, importin β1 constitutively coimmunoprecipitated with NPCs, and anti-CD3 plus anti-CD28 costimulation significantly increased their association; in contrast, in *Prkcq^-/-^* T cells, importin β1 barely bound to NPCs, regardless of stimulation (Figure 1I, Figure 1—figure supplement 1I). Similar results were found when we knocked down PKC-θ expression in Jurkat T cells with a specific small interfering RNA or short hairpin RNA (siPKC-θ or shPKC-θ, respectively) (Figure 1J-L, Figure 1—figure supplement 1J-M). These data indicate that PKC-θ deficiency alters both the basal state and TCR-induced nucleo-cytoplasmic transport.

### PKC-θ binds to RanGAP1 at the NE in a sumoylation-dependent manner

To further investigate how PKC-θ regulates nuclear transport, we constructed GST-tagged proteins including nucleoporins, RanGAP1 and other NE proteins, and performed GST pull-down assays to determine if these proteins can interact with PKC-θ. Among them, GST-RanGAP1 showed the most obvious association and a direct interaction with PKC-θ in unstimulated Jurkat cells lysate (Figure 2A,B, Figure 2—figure supplement 1A). Reciprocal co-IP from Jurkat T cells showed that PKC-θ weakly interacted constitutively with sumoylated RanGAP1, and costimulation with anti-CD3 plus anti-CD28 Abs markedly increased the association of PKC-θ with both forms of RanGAP1 (Figure 2C). A significant colocalization of PKC-θ and RanGAP1 in NPCs was also observed in Jurkat T cells after anti-CD3 plus anti-CD28 or PMA plus Iono stimulation (Figure 2—figure supplement 1B,C).

**Figure 2.**
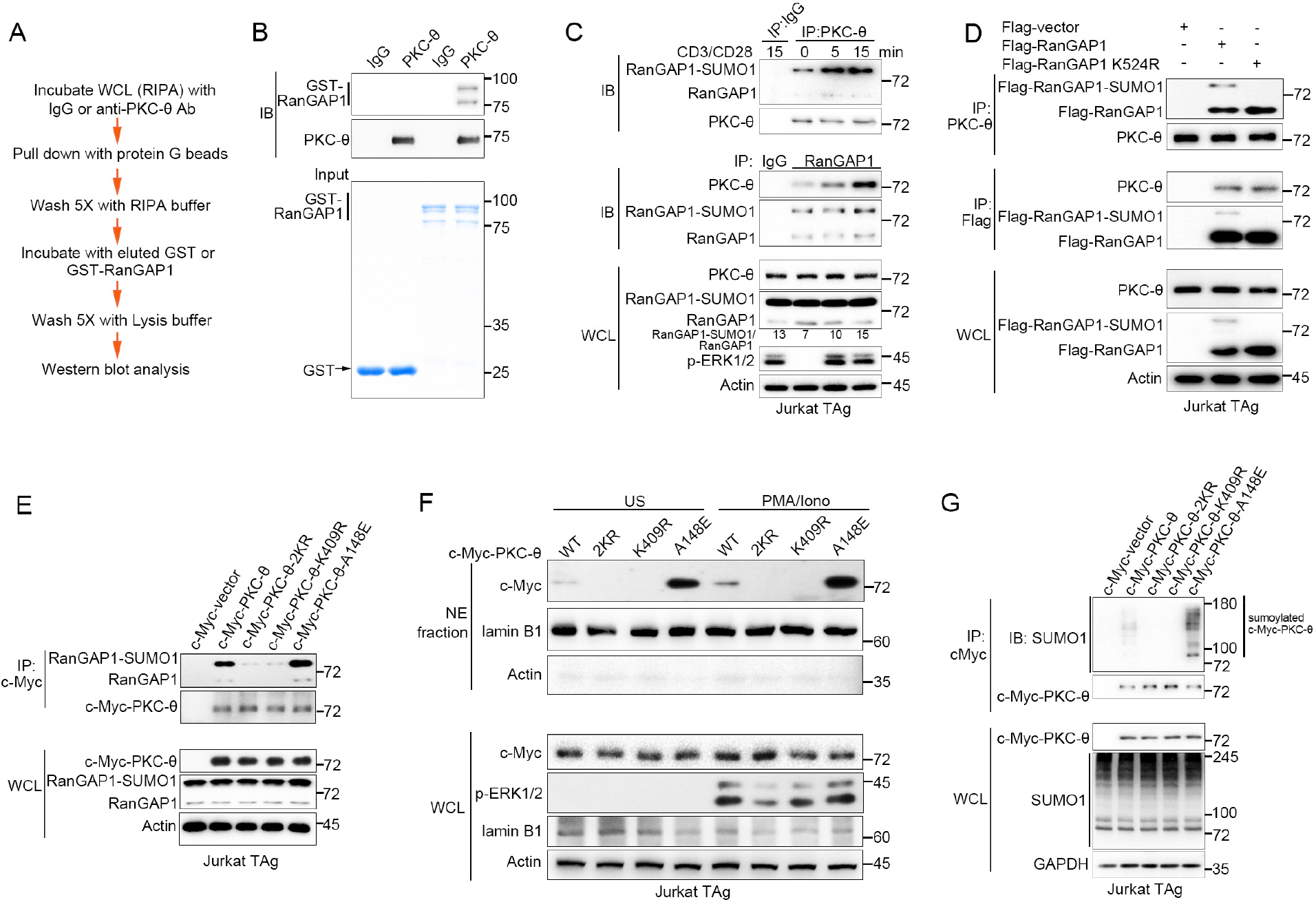
PKC-θ association with both RanGAP1 and RanGAP1-SUMO1 requires PKC-θ sumoylation. (**A**) Scheme for *in vitro* protein direct binding assay. (**B**) Analysis of direct association between RanGAP1 and PKC-θ using approach in (A). GST-fusion proteins were detected by Coomassie blue staining (bottom panel). (**C**) Immunoblot analysis of PKC-θ or RanGAP1 IPs or WCL from Jurkat E6.1 cells stimulated for 0-15 min with anti-CD3 plus anti-CD28. (**D**) Reciprocal IP analysis of the association between endogenous PKC-θ and transfected Flag-tagged wild-type or mutated RanGAP1 in Jurkat-TAg cells. (**E**) Immunoblot analysis of c-Myc-tagged PKC-θ IPs or WCL from Jurkat-TAg cells that were transiently transfected with wild-type PKC-θ or the indicated PKC-θ mutants. (**F**) Subcellular fractionation of Jurkat-TAg cells transiently transfected with wild-type PKC-θ or the indicated PKC-θ mutants (2KR, K325R/K506R desumoylation mutant; K409R, kinase dead; A148E, constitutive active), followed by immunodetection with the indicated antibodies. (**G**) Immunoblot analysis of c-Myc-tagged PKC-θ IPs or WCL from Jurkat-TAg cells that were transiently transfected with the indicated expression vectors. Data are representative of at least three biological replicates (**B-G**). **Figure supplement 1.** PKC-θ binds to and colocalizes with RanGAP1 in NE.

To elucidate the contribution of sumoylation to the association of RanGAP1 with PKC-θ, we transfected Jurkat T cells with wild-type or non-sumoylated RanGAP1 mutant (RanGAP1-K524R) having an N-terminal Flag tag or a C-terminal HA tag. PKC-θ coimmunoprecipitated efficiently with RanGAP1-K524R (Figure 2D, Figure 2—figure supplement 1D), implying that RanGAP1 sumoylation is not required for its interaction with PKC-θ. Next, we mapped the PKC-θ determinants required for its NE translocation and RanGAP1 interaction by transfecting Jurkat T cells with c-Myc-tagged wild-type PKC-θ, a desumoylated mutant PKC-θ-2KR (K325R/K506R)(Wang et al., 2015), a constitutively active PKC-θ-A148E mutant, or a catalytically inactive PKC-θ-K409R mutant. Reciprocal IP showed that both the sumoylated and non-sumoylated forms of RanGAP1 interacted more strongly with PKC-θ-A148E than with wild-type PKC-θ, whereas PKC-θ-2KR and PKC-θ-K409R displayed a much weaker interaction, if at all (Figure 2E, Figure 2—figure supplement 1E,F). Following biochemical isolation of the NE, we observed that wild-type PKC-θ translocated to the NE in response to PMA plus Iono stimulation (Figure 2F, Figure 2—figure supplement 1G), consistent with the results in Figure 1. Interestingly, PKC-θ-A148E was constitutively localized to the NE regardless of stimulation, while no apparent NE localization of either PKC-θ-2KR or PKC-θ-K409R was observed even after PMA plus Iono stimulation (Figure 2F, Figure 2—figure supplement 1G), consistent with the result in Figure 2E.

We next compared the sumoylation of wild-type PKC-θ and its mutants. As expected, wild-type PKC-θ, but not PKC-θ-2KR, was sumoylated (Figure 2G). Interestingly, PKC-θ-K409R, like PKC-θ-2KR, could not be sumoylated, whereas PKC-θ-A148E was sumoylated more strongly than wild-type PKC-θ (Figure 2G, Figure 2—figure supplement 1H), indicating that the catalytic activity of PKC-θ is required for its sumoylation. Combining the results in Figure 2E-G with the result in Figure 2D, we conclude that PKC-θ sumoylation, rather than the sumoylation of RanGAP1, was important for their association, and that PKC-θ sumoylation is required its NE translocation.

### PKC-θ deficiency inhibits the association of RanGAP1 with the NPC by reducing its sumoylation

Given the association between PKC-θ and RanGAP1, we next explored the physiological significance of this interaction. We first inspected the localization of RanGAP1 in resting T cells from wild type or *Prkcq*^-/-^ mice. Confocal microscopy revealed that NE-localized RanGAP1 was decreased in *Prkcq*^-/-^ T cells, with most RanGAP1 found in the cytosol (Figure 3A). When we used siPKC-θ to transiently knock down endogenous PKC-θ in Jurkat T cells, the amount of cytosol-localized RanGAP1 increased, while NE-localized RanGAP1 decreased (Figure 3B). Consistent with this result, stimulation of human primary T cells with anti-CD3 plus anti-CD28 or with PMA plus Iono increased the ratio of sumoylated to unsumoylated RanGAP1 (Figure 3C). A similar increase in this ratio was observed when mouse primary T cells or Jurkat T cells were costimulated with anti-CD3 plus anti-CD28 Abs; however, this increase was not observed when expression of PKC-θ was knocked out or knocked down (Figure 3D,E, Figure 3—figure supplement 1A, respectively; WCL). Consistently, PKC-θ deficiency also decreased the ratio in resting cells (Figure 3D,E, respectively; WCL and Figure 3—figure supplement 1B).

**Figure 3.**
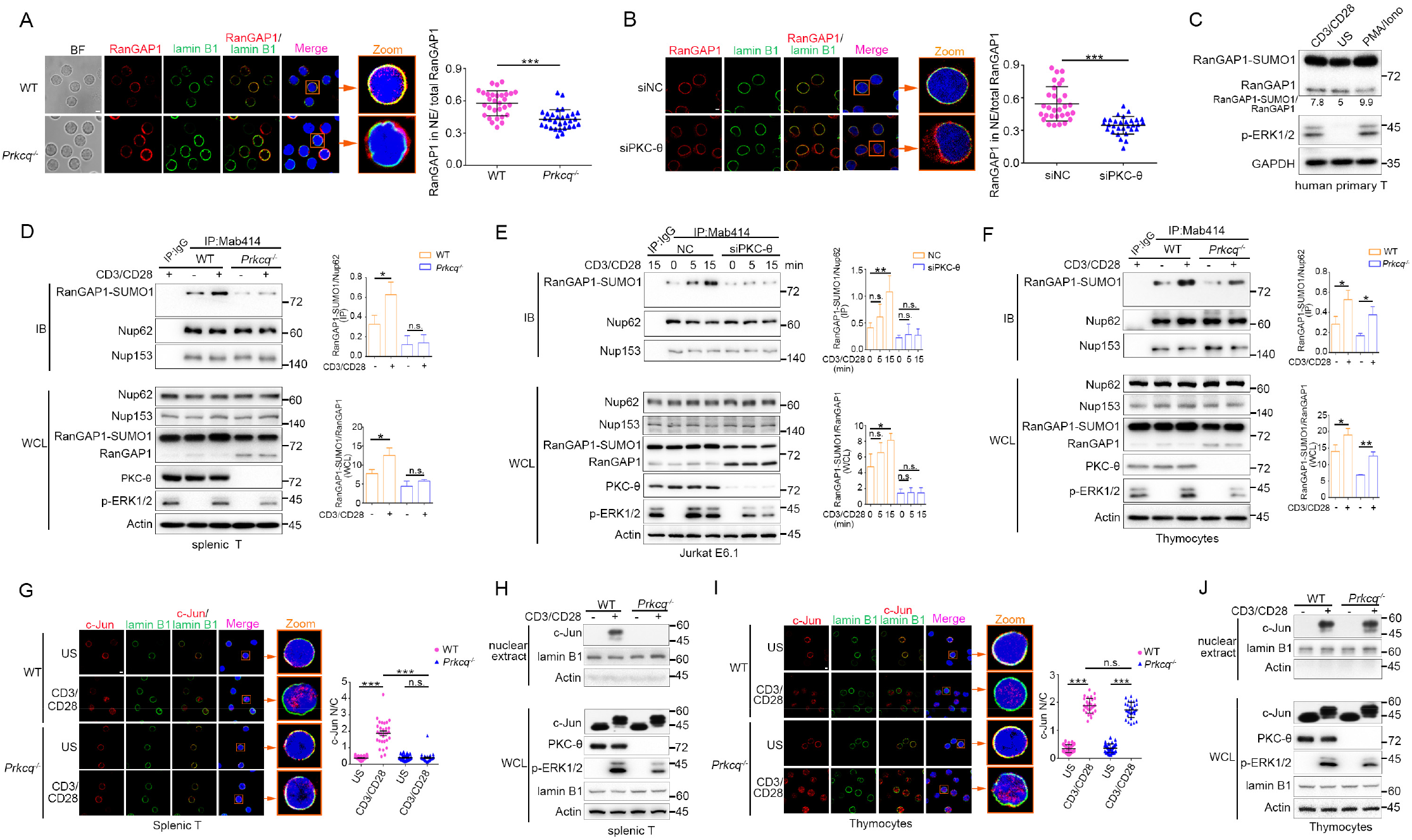
PKC-θ deficiency inhibits RanGAP1 sumoylation and its incorporation into the NPC and TCR-induced nuclear import of c-Jun in mature T cells. (**A, B**) Confocal imaging of RanGAP1 (red) and lamin B1 (green) localization in representative unstimulated WT or *Prkcq*^-/-^ mouse primary T cells (A), or in representative unstimulated Jurkat E6.1 cells transfected with siNC or siPKC-θ (B). Nuclei are stained with DAPI (blue). Areas outlined by squares in the merged images from (A) or (B) are enlarged at right in (A) or (B). Scale bars, 2 μm. Quantification of the ratio of NE/total RanGAP1 based on analysis of *~30* cells in about 6 random fields from two biological replicates as presented in (A) or (B) is shown at far right in (A) or (B), with each symbol representing an individual cell. Horizontal lines indicate the mean ± s.e.m.. \*\*\**P* < 0.001 (two tailed, unpaired Student’s t-test). (**C**) Immunoblot analysis of RanGAP1-SUMO1 and RanGAP1 in human primary T cells unstimulated or stimulated for 15 min with anti-CD3 plus anti-CD28 or PMA plus Iono. (**D-F**) Immunoblot analysis of Mab414 IPs or WCL from WT and *Prkcq*^-/-^ mouse primary splenic T cells (D) and thymocytes (F) stimulated with or without anti-CD3 plus anti-CD28 for 15 min, or from Jurkat E6.1 cells transfected with siNC or siPKC-θ stimulated for 0-15 min with anti-CD3 plus anti-CD28. (**E**) Statistical analysis of the amount ratio of RanGAP1-SUMO1 to Nup62 or to RanGAP1 in IPs or in WCL from the experiment in (D), (E) or (F) is shown at right in each figure. Analysis was based on three biological replicates for each experiment. n.s., not significant, *\*P* < 0.05, *\*\*P* < 0.01 (one-way ANOVA with post hoc test). (**G, I**) Confocal imaging of c-Jun (red) localization in representative WT or *Prkcq*^-/-^ mouse splenic T cells (G) or thymocytes (I) unstimulated (US) or stimulated with anti-CD3 plus anti-CD28, costained with anti-lamin B1 (green) and DAPI (blue). Areas outlined by squares in the merged images from (G) or (I) are enlarged at right in (G) or (I). Scale bars, 2 μm. Quantitative analysis of the N/C ratio of c-Jun analyzed in ~30 cells in about 6 random fields from two biological replicates as presented in in (G) or (I) is shown at far right in (G) or (I). Each symbol represents an individual cell. Horizontal lines indicate the mean ± s.e.m. n.s., not significant; ***P < 0.001 (one-way ANOVA with post hoc test). (**H, *J***) Subcellular fractionation of WT or *Prkcq*^-/-^ mouse splenic T cells (H) or thymocytes (J) stimulated for 0-15 min with anti-CD3 plus anti-CD28, followed by immunodetection with the indicated antibodies. Data are representative of two (**A, B, G, I**) or three (**C-F, H, J**) biological replicates. **Figure supplement 1.** PKC-θ deficiency inhibits RanGAP1 sumoylation and its incorporation into the NPC, but does not affect TCR-induced phosphorylation of c-Jun.

Sumoylated RanGAP1 was present in Mab414 NPC IPs prepared from mouse primary peripheral T cells or Jurkat T cells regardless of stimulation, but was largely absent or substantially reduced when these cells were depleted of PKC-θ both in resting and activated T cells (Figure 3D,E, Figure 3—figure supplement 1A, respectively; IP and Figure 3—figure supplement 1B). Together, these results indicate that PKC-θ promotes the sumoylation of RanGAP1 as well as the incorporation of RanGAP1 into the NPC both in resting and activated cells. Interestingly, however, and in contrast to peripheral T cells or Jurkat cells, in *Prkcq*^-/-^ mouse thymocytes, TCR plus CD28 costimulation appeared to increase the association of RanGAP1 with the NPC (Figure 3F; IP) as well as its relative sumoylation (Figure 3F; WCL) in a manner that was not affected by PKC-θ deletion, although PKC-θ deficiency did decrease the ratio of RanGAP1-SUMO1 to RanGAP1 in resting cells (Figure 3F, WCL), similar to the mature T cells (Figure 3D, WCL). Collectively, these results indicate that PKC-θ controls constitutive sumoylation in both resting mature and immature T cells and promotes the sumoylation of RanGAP1 and the association of sumoylated RanGAP1 with NPCs during TCR signaling in mature, but not in immature, T cells.

### PKC-θ deficiency inhibits TCR-induced nuclear import of c-Jun in mature but not immature T cells

Given the finding that PKC-θ is required for AP-1 activation in mature T cells but not in thymocytes (Sun et al., 2000) and the difference between thymocytes and peripheral T cells observed above (Figure 3D *vs.* Figure 3F), we considered the possibility that the activation and/or nuclear import of c-Jun, which is a component of AP-1 transcription factor, may display difference in its PKC-θ dependence in mature peripheral T cells *vs*. thymocytes. Whereas PKC-θ deficiency did not affect TCR-induced phosphorylation of c-Jun (indicative of its activation) in either mouse splenic T cells or thymocytes (Figure 3—figure supplement 1C,D), confocal microscopy and nuclear fractionation showed that the TCR-induced nuclear localization of c-Jun was blocked in *Prkcq*^-/-^ splenic T cells (Figure 3G,H, Figure 3—figure supplement 1E), but remained intact in *Prkcq*^-/-^ thymocytes (Figure 3I,J, Figure 3—figure supplement 1F), suggesting that PKC-θ regulates the nuclear import of c-Jun through the NPC in mature T cells but not in thymocytes. Thus, PKC-θ may promote c-Jun nuclear import, but not its phosphorylation *per se,* suggesting that it regulates AP-1 activation primarily by controlling the function of the NPC.

### PKC-θ-mediated phosphorylation of RanGAP1 on Ser^504^ and Ser^506^ facilitates its sumoylation

Given the fact that phosphorylation of proteins often regulates their sumoylation(Hendriks et al., 2017; Hietakangas et al., 2006; Tomasi & Ramani, 2018), we next explored the possibility that PKC-θ may regulate the sumoylation of RanGAP1 via phosphorylating it. Using a mixture of phospho-Ser- and phospho-Thr-specific antibodies, we found that TCR plus CD28 costimulation increased phosphorylation of RanGAP1 in control Jurkat T cells, but not in PKC-θ knockdown cells (Figure 4A, Figure 4—figure supplement 1A). Furthermore, an *in vitro* kinase assay demonstrated that PKC-θ immunoprecipitated from Jurkat T cells efficiently phosphorylated RanGAP1 (Figure 4B), indicating that RanGAP1 is most likely a direct PKC-θ substrate. Mass spectrometry analysis of PKC-θ-phosphorylated RanGAP1 identified five potential phosphorylation serine or threonine sites on RanGAP1 (Figure 4—figure supplement 1B). Upon mutating each of these residues individually to alanine, we found that mutation both Ser^504^ and Ser^506^, but not the three other residues, reduced the phosphorylation of RanGAP1, while phosphorylation of the double mutant S504A/S506A (RanGAP1^AA^) was significantly decreased (Figure 4C,D, Figure 4—figure supplement 1C). In the *in vitro* kinase assay, we detected phosphorylation of truncated GST-RanGAP1431-587 but not the S504A/S506A double mutant (RanGAP1^AA^; Figure 4E). Sequence alignment of RanGAP1 from different species showed that S^504^ (10/11) and S^506^ (8/11) sites and three adjacent serines are conserved (Figure 4—figure supplement 1D). A mouse RanGAP1 fragment containing these conserved sites was also phosphorylated by PKC-θ in an *in vitro* kinase assay (Figure 4—figure supplement 1E). When we calculated the ratio of RanGAP1-SUMO1 to RanGAP1 in the mutant cells, we observed that this ratio was significantly decreased in cells expressing the

**Figure 4.**
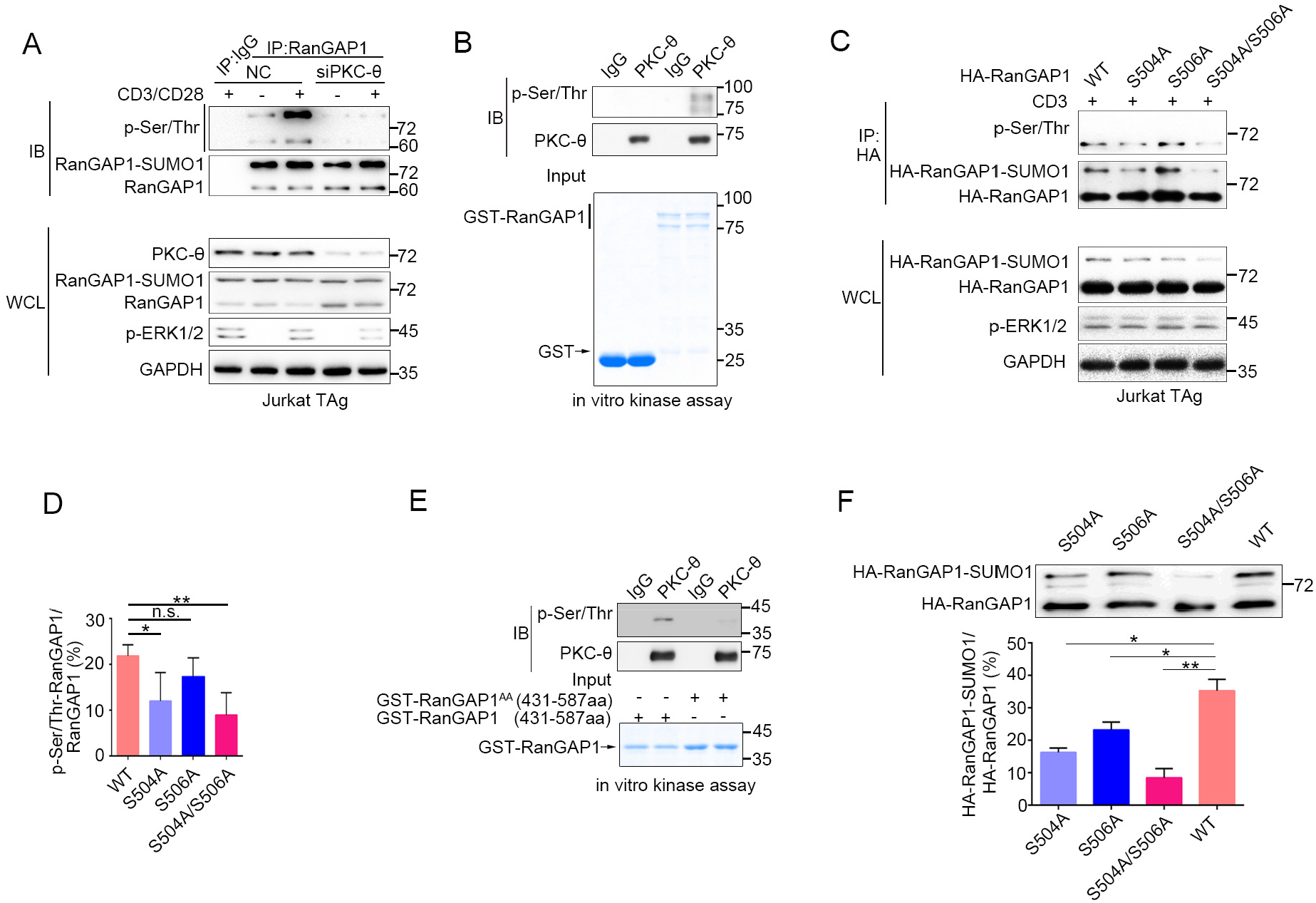
PKC-θ-mediated sumoylation of RanGAP1 requires its phosphorylation on Ser^504^ and Ser^506^. (**A**) Immunoblot analysis of serine-phosphorylated RanGAP1 in Jurkat E6.1 T cells transfected with siNC or siPKC-θ, and left unstimulated or stimulated with anti-CD3 plus anti-CD28. (**B**) *In vitro* PKC-θ kinase assay using PKC-θ immunoprecipitated from Jurkat E6.1 cells and recombinant GST-RanGAP1 as substrate. (**C, D**) Immunoblot analysis of the phosphorylation of transfected wildtype or Ser-mutated HA-tagged RanGAP1 using a mixture of p-Ser- and p-Thr-specific antibodies in Jurkat-TAg cells, which were left unstimulated or stimulated for 15 min with anti-CD3 (C). Immunoblotting of the indicated proteins in WCL is shown at bottom. The ratio of phospho-RanGAP1 to immunoprecipitated RanGAP1 is shown in (D). Analysis is based on three biological replicates. n.s., not significant, *\*P* < 0.05, *\*\*P* <0.01 (one-way ANOVA with post hoc test). (**E**) *In vitro* PKC-θ kinase assay as in (B), using purified truncated GST-RanGAP1 or GST-RanGAP1(S504A/S506A) as substrate. (**F**) Immunoblot analysis of the sumoylation of wild-type or the indicated HA-tagged RanGAP1 mutants in Jurkat-TAg cells using an anti-HA antibody (top panel). The ratios of RanGAP1-SUMO1 to RanGAP1 are shown at the bottom panel. Analysis is based on three biological replicates. n.s., not significant, *\*P* < 0.05, *\*\*P* < 0.01 (one-way ANOVA with post hoc test). Data are representative of three biological replicates. **Figure supplement 1**. PKC-θ-dependent phosphorylation of RanGAP1.

S504A mutant or the double mutant RanGAP1^AA^ and, to a lesser extent, in S506A-expressing cells (Figure 4F). Similar result was also observed when RanGAP1 antibody instead of HA antibody was used to detect HA-RanGAP1 and its SA mutants (Figure 4—figure supplement 1F). And HA-RanGAP1 and its SA mutants had a much higher expression level than endogenous RanGAP1, which could be caused by a strong promoter of the expression vector and contributed to the low sumoylation ratio of HA-RanGAP1. Together, these findings suggest that PKC-θ-mediated phosphorylation of RanGAP1, particularly on Ser^504^, promotes its sumoylation.

### PKC-θ-mediated phosphorylation of RanGAP1 is required for RanBP2/RanGAP1-SUMO1/Ubc9 subcomplex assembly

The SUMO E2 enzyme Ubc9 directly interacts with and conjugates SUMO1 to RanGAP1(Bernier-Villamor et al., 2002), and the sumoylation of RanGAP1 is essential for assembly of the RanBP2/RanGAP1-SUMO1/Ubc9 subcomplex(Hampoelz, Andres-Pons, et al., 2019; Hampoelz, Schwarz, et al., 2019; Hutten et al., 2008; Joseph et al., 2004; Mahajan et al., 1997; Reverter & Lima, 2005; Ritterhoff et al., 2016; von Appen et al., 2015; Werner et al., 2012). We therefore hypothesized that PKC-θ-mediated phosphorylation of RanGAP1 might regulate the interaction between Ubc9 and RanGAP1 and, furthermore, that RanGAP1 phosphorylation would be required for assembly of this subcomplex. Hence, we first examined whether RanGAP1 sumoylation affects its binding to Ubc9 and found that a non-sumoylated RanGAP1 mutant (K524R) was still capable of associating with Ubc9 (Figure 5A). Next, we generated two RanGAP1 mutants: One, which was mutated at both its sumoylation (K524R) and phosphorylation (S504A/S506A) sites (HA-RanGAP1^AA^/K524R), and another K524R mutant with a replacement of Ser^504^ and Ser^506^ PKC-θ phosphorylation sites by a glutamic acid as a phosphorylation mimic (HA-RanGAP1^EE^/K524R). We then analyzed the association of these mutants with Ubc9 by reciprocal co-IP and found that, compared with HA-RanGAP1-K524R, HA-RanGAP1^AA^-K524R showed a decreased association with Ubc9, while HA-RanGAP1^EE^-K524R had a stronger interaction (Figure 5A,B). This result indicates that RanGAP1 phosphorylation promotes its binding to Ubc9, and provides an explanation for our finding that mutation of the RanGAP1 PKC-θ phosphorylation sites inhibits its sumoylation (Figure 4F).

**Figure 5.**
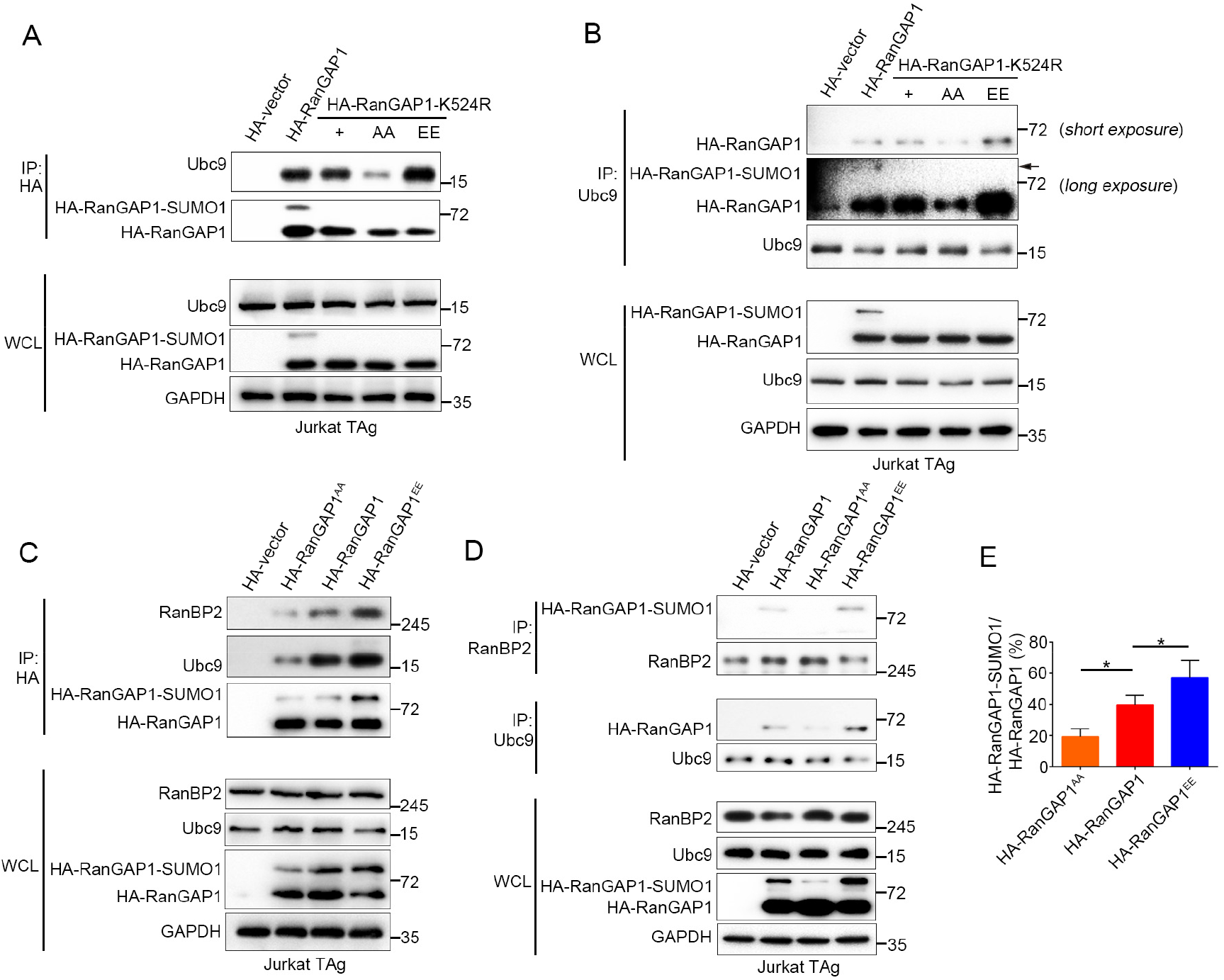
PKC-θ mediated phosphorylation of RanGAP1 is required for its association with Ubc9 and RanBP2. (**A, B**) Reciprocal IP analysis of the association between HA-tagged wild-type or K524-mutated RanGAP1 and endogenous Ubc9. RanGAP1 expression was analyzed by anti-HA antibody immunoblotting in WCL (bottom panels). (**C-E**) Immunoblot analysis of HA-RanGAP1 IPs (C) and Ubc9 IPs or RanBP2 IPs (D) from Jurkat Tag cells transfected with HA-RanGAP1 and HA-RanGAP1 mutants. The ratio of RanGAP1-SUMO1 to RanGAP1 in the WCL of (C) and (D) is quantified in (E), quantification is based on three biological replicates. *\*P* < 0.05 (one-way ANOVA with post hoc test). Data are representative of three biological replicates. **Figure supplement 1.** Characterization of the effect of Ser/Glu mutation on RanGAP1.

Next, we transfected Jurkat T cells with HA-RanGAP 1, HA-RanGAP1^AA^, or HA-RanGAP1^EE^ and analyzed their association with Ubc9 and RanBP2 by reciprocal co-IP. HA-RanGAP1^AA^ bound to Ubc9 and RanBP2 less effectively than non-mutated HA-RanGAP1; in contrast, HA-RanGAP1^EE^ bound more effectively to Ubc9 and RanBP2 (Figure 5C,D). As expected, due to its stronger association with Ubc9, the RanGAP1^EE^ mutant protein displayed an increased ratio of RanGAP1-SUMO1 to RanGAP1 relative to the two other RanGAP1 proteins (Figure 5E). Similar result was also observed when HA-RanGAP1^EE^ transfected cell lysates was blotted with RanGAP1 antibody instead of HA antibody (Figure 5—figure supplement 1A). These results reveal that PKC-θ-mediated phosphorylation of RanGAP1 is essential for assembly of the RanBP2/RanGAP1-SUMO1/Ubc9 subcomplex in NPCs. Intriguingly, *in silico* analysis showed that replacement of the two PKC-θ phosphorylation sites in RanGAP1 by glutamic acid (S504E/S506E) increased the overall structural stability of RanGAP1 (Figure 5 — figure supplement 1B); this increased stability likely contributed to the observed increased association between RanGAP1 and Ubc9. Collectively, these findings demonstrate that PKC-θ phosphorylates RanGAP1 to increase its association with Ubc9 and, in turn, the sumoylation of RanGAP1, thereby facilitating assembly of the RanBP2/RanGAP1-SUMO1/Ubc9 subcomplex upon TCR stimulation.

### RanGAP1^AA^ mutant inhibits TCR/CD28-induced AP-1, NF-ATc1 and NF-κB nuclear import and IL-2 production

Based on the results above (Figure 4 and Figure 5), we hypothesized that the non-phosphorylatable RanGAP1 mutant (RanGAP1^AA^) will have impaired ability to promote nuclear transport of key TCR-activated transcription factors that are required for productive T cell activation. As a positive control, we generated a RanGAP1 knockdown Jurkat E6.1 cell line (RanGAP1-KD) having a RanGAP1 mutation with decreased RanGAP1 protein level due to an in-frame nucleotide deletion (Figure 6—figure supplement 1A,B). We confirmed that expression of this mutant resulted in blocked TCR-induced nuclear translocation of NFATc1, p65 (NF-κB), and c-Jun and c-Fos (AP-1) (Figure 6—figure supplement 1C,D).

We next determined whether transfection of the RanGAP1-KD cell line with wild-type RanGAP1 or RanGAP1^AA^ can rescue the nuclear translocation of NFATc1, p65, or AP-1, with unedited Jurkat cells serving as a negative control. Using subcellular fractionation (Figure 6A) or confocal microscopy (Figure 6B,C), we observed that the defective nuclear translocation of these transcription factors in RanGAP1-KD cells was largely rescued by wild-type RanGAP1, but not by the RanGAP1^AA^ mutant. Similarly, while stimulated RanGAP1-KD cells displayed reduced IL-2 production, which is known to require the cooperative activity of the above transcription factors, as compared to control cells, transfection with wild-type RanGAP1 rescued IL-2 production, while RanGAP1^AA^ did not (Figure 6D). Moreover, by knocking down endogenous RanGAP1 with two different siRNAs designed to specifically target the 3’-UTR of RanGAP1 (Figure 6—figure supplement 1E), we validated the effect of the RanGAP1-AA mutation on TCR-induced nuclear translocation of the three transcription factors in human primary T cells (Figure 6—figure supplement 1E-G).

**Figure 6.**
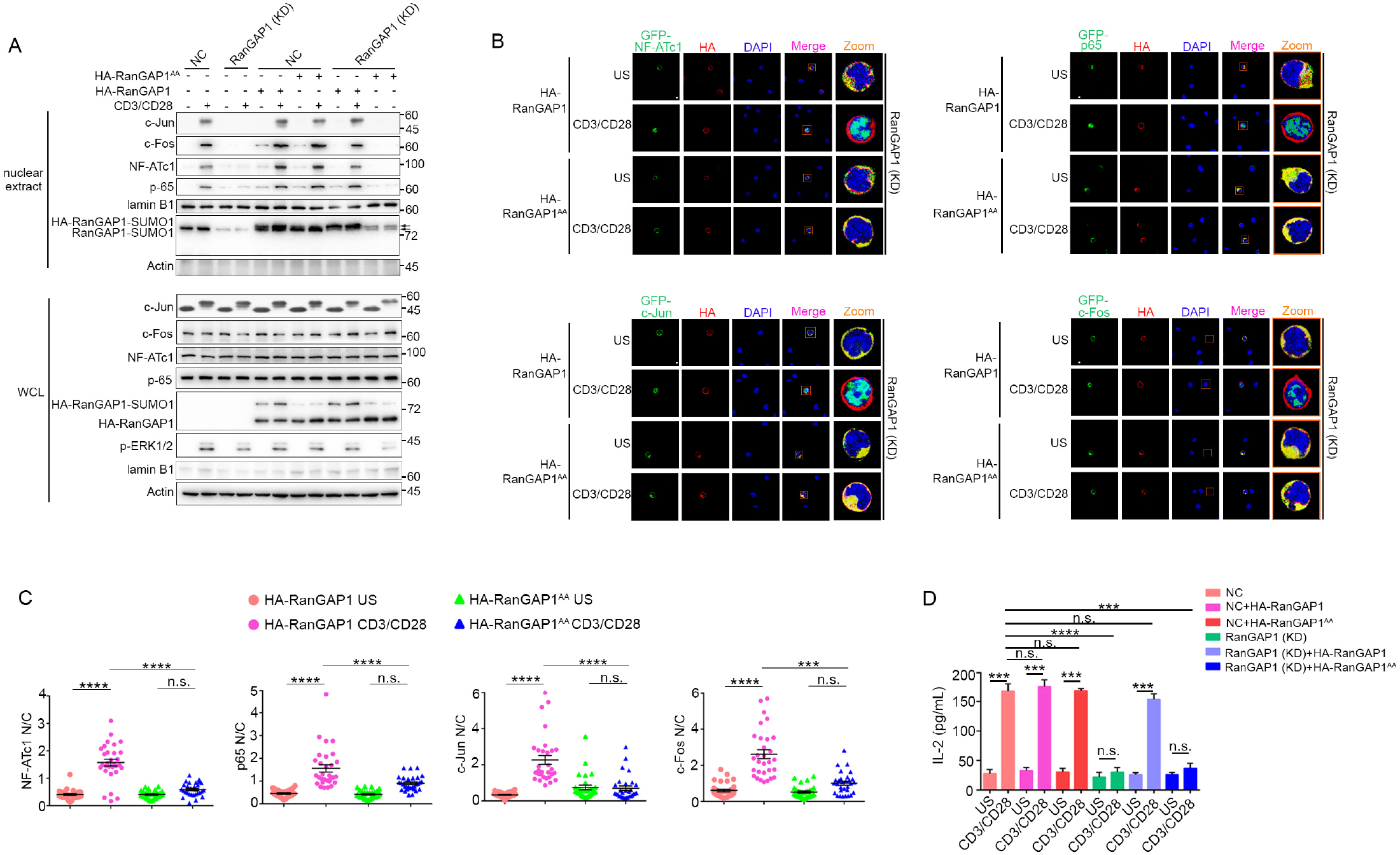
Wild-type RanGAP1, but not RanGAP1^AA^, rescues deficient TCR-induced nuclear import of NF-ATc1, NF-κB, AP-1 in RanGAP1 (KD) cells. (**A**) Immunoblot analysis of NC (cells transfected with scrambled gRNA) and RanGAP1 (KD) cells transfected with HA-RanGAP1 or HA-RanGAP1^AA^, which were left unstimulated or stimulated for 15 min with anti-CD3 plus anti-CD28, followed by nuclear fractionation and immunodetection with the indicated antibodies. (**B**) Confocal imaging of the nuclear import of GFP-tagged NF-ATc1, p65 (NF-κB), c-Jun and c-Fos in RanGAP1 (KD) cells cotransfected with HA-RanGAP1 or HA-RanGAP1^AA^, and left unstimulated or stimulated with anti-CD3 plus anti-CD28. Scale bars, 3 μm. (**C**) Quantification of the nuclear import of GFP-tagged NF-ATc1, p65 (NF-κB), c-Jun and c-Fos in ~30 cells from two biological replicates as presented in (*B*). Each symbol represents an individual T cell. Horizontal lines indicate the mean ± s.e.m. n.s., not significant, \*\*\**P* < 0.001, \*\*\*\**P* < 0.0001 (one-way ANOVA with post hoc test). (**D**) Enzyme-linked immunosorbent assay of IL-2 in supernatants of NC or RanGAP1 (KD) cells transfected with HA-RanGAP1 or HA-RanGAP1^AA^ and left unstimulated or stimulated for 24 h with anti-CD3 plus anti-CD28. n.s., not significant, \*\**P* < 0.01, *** *P* < 0.001 (one-way ANOVA with post hoc test). Data are representative of three (**A, D**) or two (**B, C**) biological replicates. **Figure supplement 1.** RanGAP1 is required for TCR-induced nuclear import of NF-ATc1, NF-κB, and AP-1.

## Discussion

PKC-θ plays an indispensable role in T cell activation, including the TCR-induced activation of NF-κB, AP-1 and NFAT, the main transcription factors required for acquisition of effector functions and cytokine production of T cells(Altman & Kong, 2016; Pfeifhofer et al., 2003; Sun et al., 2000). These PKC-θ-dependent functions depend upon its recruitment to the T cell IS and its association with CD28(Kong et al., 2011). However, little is known about potential nuclear functions of PKC-θ, with the exception of a few studies that demonstrated a role for nuclear and chromatin-associated PKC-θ in promoting gene expression(Li et al., 2016; Sutcliffe et al., 2011). Here, we have demonstrated that upon TCR plus CD28 costimulation, PKC-θ phosphorylates RanGAP1 to promote its interaction with Ubc9 and increase the sumoylation of RanGAP1, which, in turn, enhances assembly of the RanBP2 subcomplex and, thus, directly promotes the nuclear import of AP-1, NFAT and NF-κB (Figure 7). Thus, our work reveals a novel signaling axis, TCR-PKC-θ-RanGAP1, which regulates T-cell activation via control of nucleo-cytoplasmic transport, thereby linking TCR signaling to formation of the NPC.

**Figure 7.**
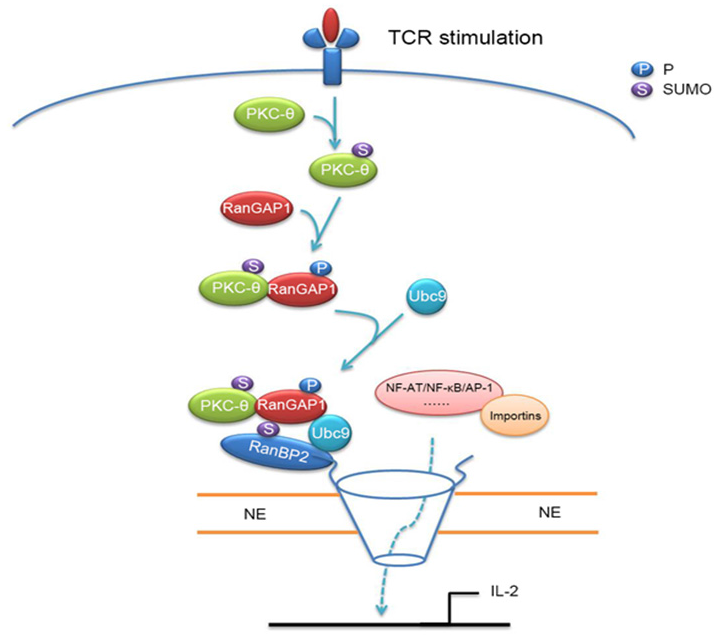
Schematic model of nuclear import regulation by the TCR-PKC-θ-RanGAP1 axis. Upon TCR stimulation, PKC-θ phosphorylates RanGAP1 to increase its association with Ubc9, thereby enhancing the sumoylation of RanGAP1, which is required for assembly of the RanBP2/RanGAP1-SUMO1/Ubc9 subcomplex. This complex then promotes the nuclear import of NF-ATc1, NF-κB, and AP-1.

Evidence for crosstalk between NPC components and immunological signaling is emerging(Borlido et al., 2018; Gu et al., 2016; Liu et al., 2015), yet much remains unknown. While the sumoylation of RanGAP1 is essential for the assembly and function of the RanBP2 complex(Matunis, Wu, & Blobel, 1998; Saitoh, Pu, Cavenagh, & Dasso, 1997), the signals that regulate this sumoylation are unclear. The sumoylation of target proteins is generally highly transient, but RanGAP1 is an exception in that it is constitutively sumoylated(Mahajan et al., 1997; Saitoh et al., 1997), a result of Ubc9 directly recognizing and catalyzing sumoylation of the RanGAP1 φ-K-x-D/E consensus motif at amino acid residues 525-528(Bernier-Villamor et al., 2002; Lee et al., 1998; Zhu, Zhang, & Matunis, 2006). The inter-regulation between phosphorylation and sumoylation is a common mechanism for the regulation of protein function(Hietakangas et al., 2006; Tomasi & Ramani, 2018). Several Ser/Thr phosphorylation sites have been identified in RanGAP1, including the phosphorylation of Ser^358^ by casein kinase II kinase to promote RanGAP1 binding to Ran protein(Takeda, Hieda, Katahira, & Yoneda, 2005), phosphorylation of Thr^409^, which is related to the nuclear accumulation of cyclin B1(Swaminathan et al., 2004), and mitotic phosphorylation of Ser^428^ and Ser^442^ with unknown function(Swaminathan et al., 2004). However, none of these phosphorylations affect the sumoylation of RanGAP 1. In contrast, phosphorylation of Ser^504^ and Ser^506^ by PKC-θ, which we documented here, promoted the sumoylation of RanGAP1 by enhancing the interaction between RanGAP1 and Ubc9, which may be facilitated by these two phosphorylation sites being close to the φ-K-x-D/E motif. Thus, our study suggests a unique role for PKC-θ in linking TCR signaling to assembly of the RanBP2 complex.

Our finding that PKC-θ deficiency essentially abolished the binding of RanGAP1-SUMO1 to NPC while only moderately inhibiting the sumoylation of RanGAP1 upon stimulation *in vivo* (Figure 3D,E) is of interest. A possible explanation for this apparent discrepancy is that the increased RanGAP1 phosphorylation itself, mediated by PKC-θ upon stimulation, may also assist the binding. Moreover, other PKC-θ substrates in addition to RanGAP1 may exist in the NPC complex, thus, phosphorylation of these proteins by PKC-θ may contribute to the TCR-induced NPC assembly as well. The T cell adaptor protein, SLP76, has been reported to bind to RanGAP1-SUMO1 and to promote the TCR-induced nuclear import of NFAT(Liu et al., 2015). Combining these findings, we propose that some TCR signaling modules, *e.g.,* PKC-θ and SLP76, translocate to the NE to regulate nucleo-cytoplasmic transport and, furthermore, that PKC-θ plays an indispensable role in NPC assembly.

The functions of PKC-θ in T cell activation and immunity are largely determined by its cellular localization in the central region of the IS and by its association with CD28, two requirements for productive T cell activation(Kong et al., 2011; Wang et al., 2015; Yokosuka et al., 2008). In the IS, PKC-θ phosphorylates CARMA1 to facilitate the formation of CARMA1-Bcl10-MALT complex and, in turn, activates the canonical IKK-NF-κB signaling pathway(Matsumoto et al., 2005; Ruland, Duncan, Wakeham, & Mak, 2003). Here, we demonstrate another, rather indirect and previously unknown role of PKC-θ in mediating NF-κB activation, namely, enabling the nuclear import of NF-κB via promoting sumoylation of RanGAP1 and, consequently, assembly of the RanBP2 subcomplex. Our findings reveal that this action of PKC-θ also facilitates the nuclear import of two other transcription factors required for productive T cell activation, *i.e.*, NFATc1 and AP-1.

Upon TCR stimulation, PKC-θ also translocates to the nucleus, where it associates with chromatin and forms transcription complexes with Pol II, MSK-1, LSD1, and 14-3-3ζ (Sutcliffe et al., 2011) or phosphorylates a key splicing factor, SC35, and histone (Li et al., 2016; McCuaig et al., 2015) to activate downstream gene transcription. Here, we show that sumoylated PKC-θ translocates to the NE to promote the function of the NPC. Interestingly, while the nuclear localization sequence of PKC-θ is known to mediate its nuclear localization(Sutcliffe et al., 2012), desumoylation of PKC-θ or inactivation of its catalytic activity prevented its NE localization (Figure 2F). Thus, different mechanisms may be involved in PKC-θ translocation to the NE *vs.* the nucleus. It is, therefore, possible that localization of PKC-θ to the NE promotes the function of NPC, which, in turn, would enable the nuclear translocation of cargo PKC-θ. We have previously demonstrated that PKC-θ sumoylation, catalyzed by the SUMO E3 ligase PIASxβ, is required for its central IS localization(Wang et al., 2015). It remains to be determined whether PKC-θ translocation to the NE similarly requires or is dependent on another mechanism.

The RanBP2 subcomplex is a multifunctional component of NPC in the animal kingdom (Hampoelz, Andres-Pons, et al., 2019; D. H. Lin & Hoelz, 2019). In addition to its roles in nuclear transport and NPC assembly(Hutten et al., 2008), the RanBP2 subcomplex has other cellular functions, including translational control (Mahadevan et al., 2013) and nuclear import of pathogens(Dharan et al., 2016). Moreover, the RanBP2/RanGAP1-SUMO1/Ubc9 complex is a multi-subunit SUMO E3 ligase (Reverter & Lima, 2005; Werner et al., 2012) that mediates the sumoylation of the GTPase Ran (Hutten et al., 2008; Sakin, Richter, Hsiao, Urlaub, & Melchior, 2015; Walde et al., 2012) and the sumoylation of topoisomerase IIa and borealin to regulate mitosis(Dawlaty et al., 2008; Klein, Haindl, Nigg, & Muller, 2009). Therefore, our current findings that reveal a TCR-PKC-θ-RanGAP1 signaling axis not only exposes a novel regulatory layer of TCR signaling, but also provides a new angle to understand fundamental mechanisms of T cell immunity.

## Materials and Methods

### Mice

C57BL/6 (B6) and *Prkcq*^-/-^ mice (a gift from D. Littman) were housed under specific pathogen-free conditions and were manipulated according to guidelines approved by the Animal Care and Ethics committee of Sun Yat-Sen University.

### Plasmids

The cDNAs encoding RanGAP1, p65/NF-κB, NF-ATc1, c-Jun, c-Fos were amplified by PCR from a Jurkat E6.1 T cell cDNA library and were cloned into the vectors pcDNA3.1-HA (Invitrogen), pGEX-4T-2 (Sigma), or pcDNA3.1-GFP (Invitrogen). Plasmids encoding HA-tagged SUMO1, and c-Myc-tagged PKC-θ, PKC-θ-2KR, PKC-θ-K409R, PKC-θ-A148E have been described (Wang et al., 2015). Specific point mutations of RanGAP1 were introduced by site-directed mutagenesis with a QuikChange Site-Directed Mutagenesis Kit (Stratagene).

### Antibodies and cytokines

Antibodies specific for PKC-θ (C-19), RanGAP1 (C-5), c-Myc (9E10), HA (Y-11), actin (I-19), GAPDH (Fl-335), importin β1 (H-7), Ran (A-7), p-Ser (4A3), NF-ATc1 (4-3), p65/NF-κB (sc-109), lamin B1 (S-20), GFP/YFP (B-2), Ubc9 (C-12), and RanBP2 (D-4) were from Santa Cruz Biotechnology. Antibody to PKC-θ (610089) was from BD Biosciences. Antibodies to phospho-Ser/Thr (9631), c-Jun (60A8), c-Fos (9F6), phospho-c-Jun (9261), and phospho-ERK1/2 (T202), Na/K-ATPas (3010) were from Cell Signaling Technology. Mab414 (902901), anti-mouse CD3 (145-2C11) and anti-mouse CD28 (37.51) were from BioLegend. Anti-human CD3 (OKT3) and anti-human CD28 (CD28.2) were from eBioscience. Goat anti-mouse IgG (31160) was from Thermo Fisher Scientific. Alexa Fluor 488-coupled chicken anti-mouse IgG (A-21200), Alexa Fluor 594-coupled chicken anti-mouse IgG (A-21201), Alexa Fluor 594-coupled chicken anti-rabbit IgG (A-21442), Alexa Fluor 488-coupled donkey anti-goat IgG (A-11055), Alexa Fluor 647-coupled donkey anti-goat IgG (A-21447), and CellTrace Violet Cell Proliferation dye (C34557) were from Invitrogen.

### Human blood

De-identified human peripheral blood was obtained from healthy adult donors and was provided by the Guangzhou blood center; it was handled according to the guidelines of the Animal Care and Ethics committee of Sun Yat-Sen University. Peripheral blood mononuclear cells isolation and T cell enrichment were performed as previously described(Wang et al., 2015).

### Cell culture, transfection and stimulation

Spleens and thymii of *Prkcq*^-/-^ mice were dissociated into singlecell suspensions in PBS containing 1% FBS (Gibco), and samples were depleted of red blood cells with RBC lysis buffer (Sigma). Mouse splenic T cells were isolated with a Pan T Cell Isolation Kit II (Miltenyi Biotec). Jurkat T cells and Raji B cells were cultured in complete RPMI-1640 (Hyclone, Logan, UT, USA) medium supplemented with 10% FBS, and 100 U/ml each of penicillin and streptomycin (Life Technologies). Jurkat E6.1 T cells stably expressing PKC-θ-specific short hairpin RNA (shPKC-θ) were grown in the presence of aminoglycoside G418 (700 μg/ml; Invitrogen). Cells in a logarithmic growth phase were transfected by electroporation. For APC stimulation of T cells, Raji B lymphoma cells were incubated for 30 min at 37°C in the presence or absence of SEE (100 ng/ml; Toxin Technology). The cells were washed with PBS and were mixed with Jurkat T cells at a ratio of 1:1, followed by incubation for various times at 37°C. For antibody stimulation, mouse or human T cells were stimulated for various times with anti-CD3 (5 μg/ml) and/or anti-CD28 (2 μg/ml), which were crosslinked with goat anti-mouse IgG (10 μg/ml).

### IP and immunoblotting

Cells washed with ice-cold PBS and lysed in lysis buffer [20 mM Tris-HCl, pH 7.5, 150 mM NaCl, 5 mM EDTA, 1% Nonidet-P-40, 5 mM NaPPi, 1 mM sodium orthovanadate (Na_3_VO_4_), 1 mM PMSF, and 10 μg/ml each aprotinin and leupeptin]. Whole cell lysates were incubated overnight at 4°C with the indicated antibodies, and proteins were collected on protein G-Sepharose beads (GE Healthcare) for an additional 4 h at 4°C with gentle shaking. The immunoprecipitated proteins were resolved by SDS-PAGE, transferred onto PVDF membranes and probed with primary antibodies. Signals were visualized by enhanced chemiluminescence (ECL; YESEN, Shanghai, CHINA) and films were exposed in the ChemiDoc XRS+ system (Bio-Rad) or to X-ray film. Densitometry analysis was performed with ImageJ software.

### NE and NPC fractionation

The nuclear and NE fractions were prepared using the NE-PERTM Nuclear and Cytoplasmic Extraction kit (Thermo Scientific) as per manufacturer’s instructions. Briefly, cells washed with ice-cold PBS were resuspended in Cytoplasmic Extraction Reagents containing proteinase inhibitors, vortexed vigorously and centrifuged at 16,000 x *g* for 10 min. The pellet was resuspended in Nuclear Extraction Reagent containing proteinase inhibitors, vortexed vigorously and centrifuged at 16,000 x *g* for 10 additional min. The supernatant and the insoluble fraction, representing the nuclear extract and NE, respectively, were collected.

The NPC fraction was prepared as described(Jafferali et al., 2014), with minor modifications. Cells were washed with PBS and treated with 1 mM dithiobis (succinimidyl propionate) (DSP, Sangon Biotech) in RPMI-1640 medium for 15 min at room temperature to crosslink the NPC. The reaction was stopped by adding 15 mM Tris-HCl (pH 7.4) for 10 min at room temperature. Nuclei were pelleted as described above, followed by incubation in 5 volumes of 7 M urea containing 1% Triton X-100 (TX-100) and protease inhibitors for 20 min on ice to resuspend the nuclei pellet. The suspension was collected as the NPC fraction, diluted 8-fold in PBS containing protease inhibitors, sonicated on ice, and cleared by centrifugation at 1,000 x *g* for 10 min.

### Fluorescence microscopy and analysis

Immunofluorescence was conducted as previously described(Wang et al., 2015). Briefly, conjugates of Jurkat T cells and Raji APCS were plated on poly-l-lysine-coated slides, incubated for 15 min at room temperature, fixed for 15 min with 4% PFA, and permeabilized with 0.2% Triton X-100 for 10 min at room temperature. The slides were blocked with 2% BSA for 1 h, and samples were stained with indicated antibodies overnight at 4°C. After washing with PBS, slides incubated for 1 h at room temperature with secondary antibodies. After three washes with PBS, the cells were mounted with a drop of mounting medium. Images were obtained with a Leica SP5 laser-scanning confocal microscope equipped with 100× objective lens with laser excitation at 405 nm, 488 nm, 561 nm, or 633 nm. Each image is a single z-section and the z-position closest to the center of the cell (the equatorial plane) was chosen. Images were analyzed and processed with ImageJ and Adobe Photoshop CS6 software. Briefly, quantitative colocalization analysis of confocal microscopy images was performed with the JACoP module of the FIJI-ImageJ software (https://imagej.nih.gov/ij/index.html) to generate the Pearson’s correlation coefficient (PCC) with a range of+1 (perfect correlation) to −1 (perfect exclusion). PCC measures the pixel-by-pixel covariance in the signal levels of two channels of an image. The protein nuclear/cytoplasmic (N/C) expression ratio of confocal microscope images was quantified as follows: N/C = Total fluorescence intensity in nuclear area/(Total fluorescence intensity in whole cell area - total fluorescence intensity in nucleus). The fluorescence intensities of PKC-θ or RanGAP1 in the NE were quantified with FIJI-ImageJ software. The region of nuclear envelope was automatically segmented as described with FIJI-ImageJ software(Tosi, Bardia, Filgueira, Calon, & Colombelli, 2020).

### Expression and purification of GST-fusion proteins

GST-fusion proteins were expressed in *E. coli* BL21 after induction with 0.3 mM isopropyl-beta-D-thiogalactopyranoside (Sangon Biotech) for 12 h at 18°C. Bacteria were resuspended in lysis buffer (1XPBS: 137mM NaCl, 2.7mM KCl, 10mM Na2HPO4, and 1.8 mM KH2PO4, PH7.4; proteinase inhibitors, and 1% Triton X-100 for GST-Nups or 0.1% Triton X-100 for GST-RanGAP1). Bacterial extracts were sonicated for 10 min and centrifuged. GST-fusion proteins were purified by incubation with glutathione-Sepharose beads(GE Healthcare). The precipitates were washed 3X with lysis buffer, then eluted with elution buffer (50 mM Tris-HCL, pH 8.0, 15 mM reduced glutathione). Coomassie Brilliant Blue (CBB) staining was used as loading control.

### PKC-θ *in vitro* kinase assay

The kinase assay was conducted as previously described(Wang et al., 2015). Jurkat T cell lysates were immunoprecipitated with anti-PKC-θ or a control IgG. The IPs were washed 5X with RIPA buffer containing 0.2% SDS and 1X with PKC-θ kinase buffer (20 mM HEPES, pH 7.2-7.4, 10 mM MgCl2 and 0.1 mM EGTA), and were resuspended in 25 μl of kinase buffer containing 20 μM cold ATP, 10 μM PMA, 200 μg/mL phosphatidyserine and 1 μg of recombinant GST-RanGAP1 protein or GST-RanGAP1 mutant proteins as substrate. After incubation for 30 min at 30°C with gentle shaking, the reaction was stopped by adding 5X loading buffer. Samples were boiled at 95°C for 10 min, separated by SDS-PAGE and detected by western blot with an anti-p-Ser/Thr antibody (a mixture of phospho-Ser-and phospho-Thr-specific antibodies, BD Biosciences, 9631).

### Mass Spectrometric Analysis

Samples for co-IP were prepared as described previously with minor modification(Wang et al., 2015). In brief, anti-CD3/CD28-stimulated Jurkat T cells were lysed, and followed by IP with anti-PKC-θ or anti-IgG. IPs immobilized on protein G beads and 1 μg recombinant GST-RanGAP 1 protein immobilized on GSH Sepharose beads were separately washed 2X with kinase buffer, and mixed to initiate the kinase assay. Reactions were terminated by Laemmli sample buffer, boiled, and resolved on SDS-PAGE. Gel bands of interest were excised and subjected to tryptic digestion. After desalting, the peptides were analyzed by tandem MS. A splitless Ultra 2D Plus system (Eksigent) coupled to the TripleTOF 5600 System (AB SCIEX) with a Nanospray III source (AB SCIEX) were performed to analyze immunoprecipitated proteins and identify posttranscriptional modification sites of RanGAP1. The search engine ProteinPilot V4.5 was used to assigned potential modification sites with high confidence.

### Immunoelectron microscopy

Immunoelectron microscopy was performed on Jurkat T cells stimulated for 015 min with anti-CD3 plus anti-CD28. Cells were fixed in buffer (4% paraformaldehyde, 0.2% glutaraldehyde), pelleted, treated with LR white acrylic resin (L9774, Sigma-Aldrich), and then frozen for 72 h at −20°C with UV irradiation. Frozen pellets were sectioned by a cryo-ultramicrotome (EM UC6 and FC6, Leica). Cryosections were thawed, rinsed in PBS with 1% glycine, and incubated in 0.01 M PBS containing 0.1% BSA and 5% goat serum for 30 min at room temperature. The samples were incubated with mouse anti-PKC-θ antibody diluted 1:50 overnight at 4°C, and rinsed in 0.01 M PBS, then incubated with 10 nm colloidal gold-labeled anti-mouse IgG secondary antibody (G7652, Sigma-Aldrich) diluted 1:25 for 3 h at room temperature. Grids were then rinsed in 0.01 M PBS and ultrapure water, and embedded in 2% uranyl acetate with lead citrate. Images were taken on an electron microscope (JEM1400, JEOL).

### Enzyme-linked immunosorbent assay (ELISA)

Samples for ELISA were prepared as described previously (Wang et al., 2015). Aliquots of T cells (3 × 10^6^) transfected with HA-RanGAP1 or HA-RanGAP1^AA^ vectors were stimulated for 24 h with anti-CD3 plus anti-CD28, and the concentration of IL-2 in culture supernatants was determined by ELISA according to the manufacturer’s instructions (BD Biosciences).

### Computational Analysis

I-Mutant2.0 (folding.biofold.org/i-mutant/i-mutant2.0.html) was using to predict RanGAP1 protein stability changes with single point mutation from the protein sequence (NP_002874.1).

### Statistical analysis

Prism (GraphPad 5.0 Software) was used for graphs and statistical analysis. Statistical analysis was performed with a two-tailed, unpaired Student’s */*-test or one-way ANOVA with post hoc test. *P* values of less than 0.05 were considered statistically significant. Graphs represent mean ± standard error of the mean (s.e.m).

## ACKNOWLEDGMENTS

Supported by the National Natural Science Foundation of China (31670893, 31370886, 31170846), and the Guangzhou Science and Technology Project (201904010445).

## AUTHOR CONTRIBUTIONS

Y.L. conceived and supervised this study; Y.L. and Y.H. designed experiments, analyzed data and wrote the manuscript; Y.H., Z.Y., CS.Z., Y.G., Z.X., YY.L., Y.C. and D.F. performed experiments. A.A contributed critical reagents and suggestions, and edited the manuscript.

## Supplementary file 1 - Primers and siRNA sequences

**Figure 1—figure supplement 1.**
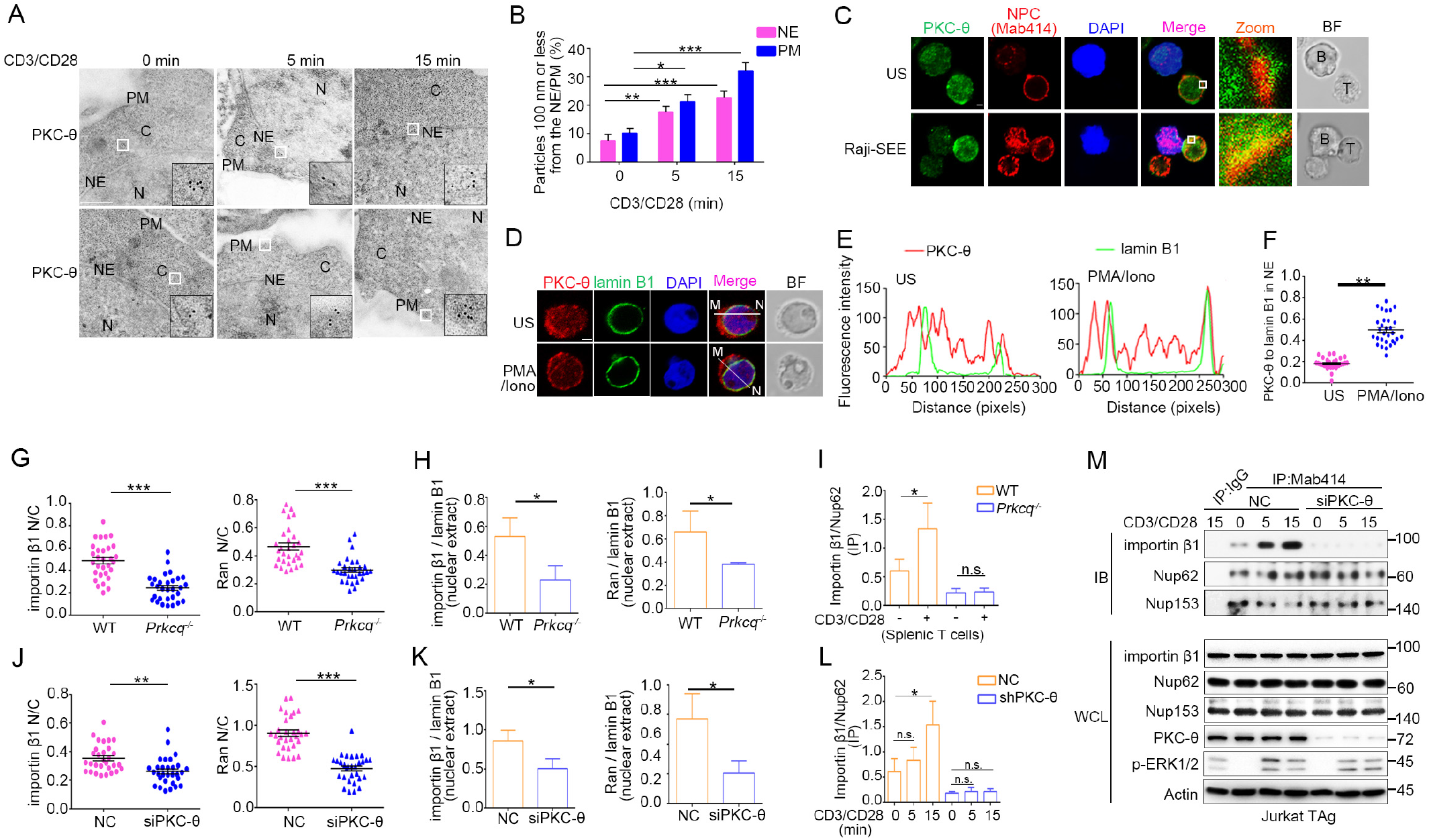
PKC-θ translocates to the NE following TCR stimulation and PKC-θ deficiency decreases nuclear import of importin β1 and Ran and NPC association with importin β1. (**A**) Transmission electron microscopy (TEM) images of PKC-θ labelled by antibody-conjugated gold particles in Jurkat E6.1 cells stimulated for 0-15 min with anti-CD3 plus anti-CD28. The areas outlined by small white squares in middle are enlarged at bottom right corner (black squares). C, cytoplasm; N, nucleus; NE, nuclear envelope; PM, plasma membrane. Scale bars, 500 nm. (**B**) Quantitation of the percentage of gold particles localized ≤100 nm from the NE or from the PM in cells from (A). *\*P* < 0.05, *\*\*P* < 0.01, *\*\*\*P* < 0.001 (one-way ANOVA with post hoc test). (**C**) Confocal imaging of PKC-θ and NPCs colocalization in Jurkat E6.1 cells unstimulated (US) or stimulated for 15 min with superantigen (SEE)-pulsed Raji B cells that have been labelled with Cell Tracker Blue (blue). Areas outlined by squares in the merged images are enlarged at right. Scale bars, 2 μm. (**D, E**) Confocal imaging showing colocalization of endogenous PKC-θ (red) with lamin B1 (green) in representative Jurkat E6.1 cells left unstimulated or stimulated for 15 min with PMA plus Iono (D). Nuclei are stained with DAPI (blue). Lamin B1 marks the NE. Scale bars, 2 μm. Pixel intensity along a line from M to N in the merged images is shown in (E). (**F**) Quantification of ~30 cells with NE-localized PKC-θ from two biological replicates in (D). Each symbol represents an individual T cell. Horizontal lines indicate the mean ± s.e.m. *\*\*P* < 0.01 (two tailed unpaired Student’s Λtest). (**G**) Quantification of the N/C ratio of importin β1 (left) and Ran (right) based on analysis of ~30 cells in about 6 random fields from two biological replicates of Figure 1G. Horizontal lines indicate the mean ± s.e.m. *\*\*\*P* < 0.001 (two tailed unpaired Student’s Λtest). (**H**) Statistical analysis of importin β1 and Ran in nuclear extract from the experiment of Figure 1H. Analysis was based on three biological replicates. *\*P* < 0.05 (one-way ANOVA with post hoc test). (**I**) Statistical analysis of importin β1 binding to NPCs in Mab414 antibody IPs from the experiment of Figure 1I. Analysis was based on three biological replicates. n.s., not significant, *\*P* < 0.05 (oneway ANOVA with post hoc test). (**J**) Quantification of the N/C ratios of importin β1 (left) and Ran (right) based on analysis of *~*30 cells in about 6 random fields from two biological replicates of Figure 1J. Horizontal lines indicate the mean ± s.e.m. *\*\*P* < 0.01, *\*\*\*P* < 0.001 (two tailed unpaired Student’s Λtest). (**K**) Statistical analysis of importin β1 and Ran in nuclear extract from the experiment of Figure 1K. Analysis as based on three biological replicates. *\*P* < 0.05 (one-way ANOVA with post hoc test). (**L**) Statistical analysis of importin β1 binding to NPCs (immunoprecipitated with Mab414 antibody) in the experiment shown in Figure 1L. Analysis was based on three biological replicates. n.s., not significant, *\*P* < 0.05, *\*\*P* < 0.01 (one-way ANOVA with post hoc test). (**M**) Immunoblot analysis of NPC IPs (Mab414) or WCL from unstimulated or stimulated Jurkat E6.1 T cells transfected with scrambled siRNA negative control (siNC) or PKC-θ targeting siRNA (siPKC-θ). Data are representative of two (**A-F**) or three (**G-M**) biological replicates.

**Figure 2—figure supplement 1.**
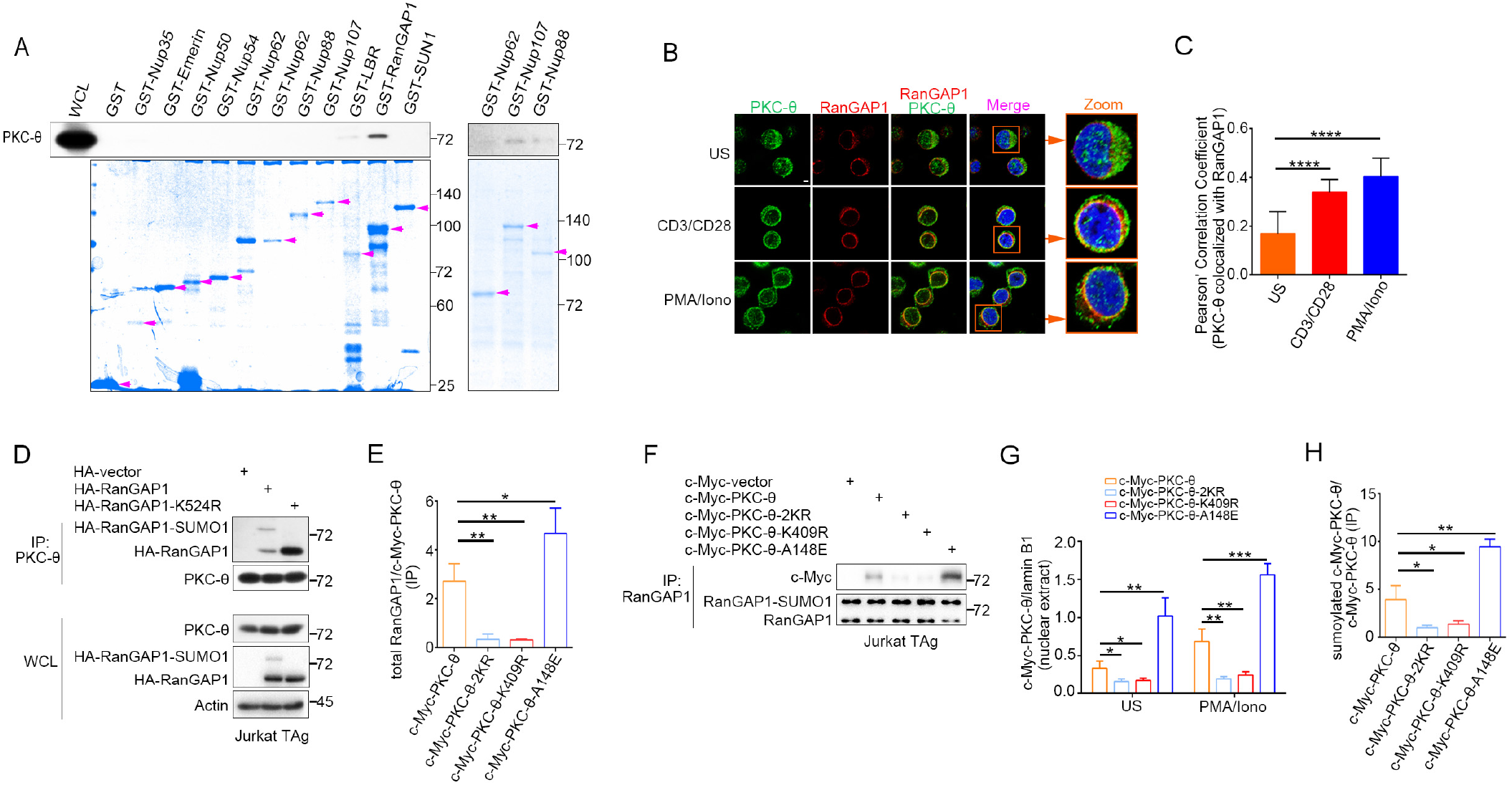
PKC-θ binds to and colocalizes with RanGAP1 in NE. (**A**) Association of PKC-θ with the indicated GST fusion proteins representing different NPC components demonstrated by a pull-down assay from unstimulated Jurkat E6.1 cell. The recombinant proteins were revealed by Coomassie blue staining. Purple arrows indicate the bands corresponding to GST, GST-fusion proteins. (**B**) Confocal imaging of intracellular of PKC-θ (green) and RanGAP1(red) colocalization in Jurkat E6.1 cells left unstimulated or stimulated for 15 min with anti-CD3 plus anti-CD28 or with PMA plus Iono. Cell nuclei are stained with DAPI. Areas outlined by squares in the merged images are enlarged at right. Scale bar, 2 μm. (**C**) Quantitative analysis of colocalization of PKC-θ with RanGAP1 shown in (B). Statistical analysis was based on at least three different colocalization images covering dozens of cells using the ImageJ software. (**D**) Immunoblot analysis of endogenous PKC-θ IPs from Jurkat-TAg cells, detected by HA antibody. (**E**) Statistical analysis of RanGAP1 binding to PKC-θ in c-Myc antibody IPs from the experiment of Figure 2E. Analysis was based on three biological replicates. *\*P* < 0.05, *\*\*P* < 0.01 (one-way ANOVA with post hoc test). Total RanGAP1 included RanGAP1 and RanGAP1-SUMO1. (**F**) Immunoblot analysis of endogenous RanGAP1 IPs from Jurkat-TAg cells transfected with the indicated c-Myc-tagged PKC-θ expression vectors. Data are representative of three independent experiments. (**G**) Statistical analysis of c-Myc-PKC-θ in nuclear extract from the experiment of Figure 2F. Analysis was based on three biological replicates. *\*P* < 0.05, \*\**P* < 0.01, \*\*\**P* < 0.001 (two-way ANOVA). (**H**) Statistical analysis of c-Myc-PKC-θ sumoylation from the experiment of Figure 2G. Analysis was based on three biological replicates. *\*P* < 0.05, *\*\*P* < 0.01(one-way ANOVA with post hoc test). Data are representative of three biological replicates.

**Figure 3—figure supplement 1.**
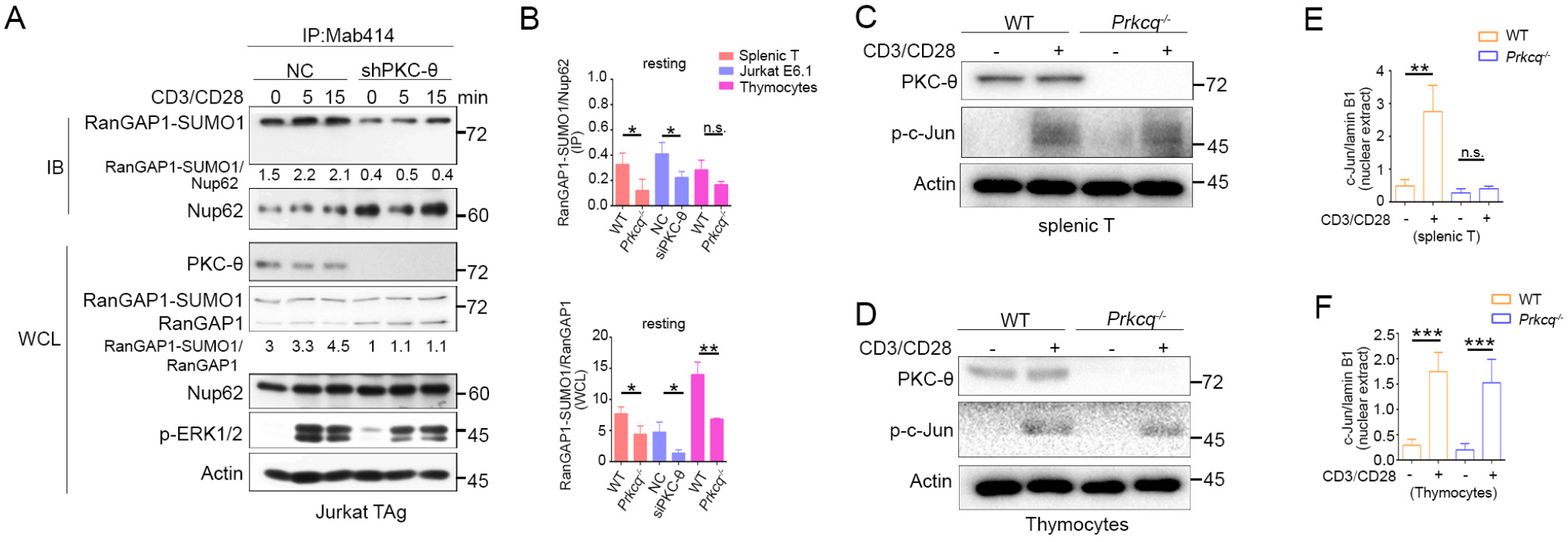
PKC-θ deficiency inhibits RanGAP1 sumoylation and its incorporation into the NPC, but does not affect TCR-induced phosphorylation of c-Jun. (**A**) Immunoblot analysis of the association between RanGAP1-SUMO1 and NPC in Mab414 IPs, and of the indicated proteins expression in WCL from NC and shPKC-θ Jurkat T cells stimulated for 0-15 min with anti-CD3 plus anti-CD28. (**B**) Statistical analysis of the amount ratio of RanGAP1-SUMO1 to Nup62 or to RanGAP1 in IPs or in WCL under unstimulated condition from the experiment in Figure 3D-F. Analysis was based on three biological replicates. n.s., not significant, *\*P < 0.05, **P* < 0.01 (one-way ANOVA with post hoc test). (**C, D**) Immunoblot analysis of phospho-c-Jun in lysates of WT or *Prkcq*^-/-^ mouse peripheral T cells (C) or thymocytes (D) that were left unstimulated (US) or stimulated with anti-CD3 plus anti-CD28 for 15 min. (**E,F**) Statistical analysis of c-Jun nuclear import from the experiment of Figure 3H(E), J(F). Analysis was based on three biological replicates. n.s., not significant, \*\**P* < 0.01, \*\*\**P* < 0.001 (one-way ANOVA with post hoc test). Data are representative of three biological replicates.

**Figure 4—figure supplement 1.**
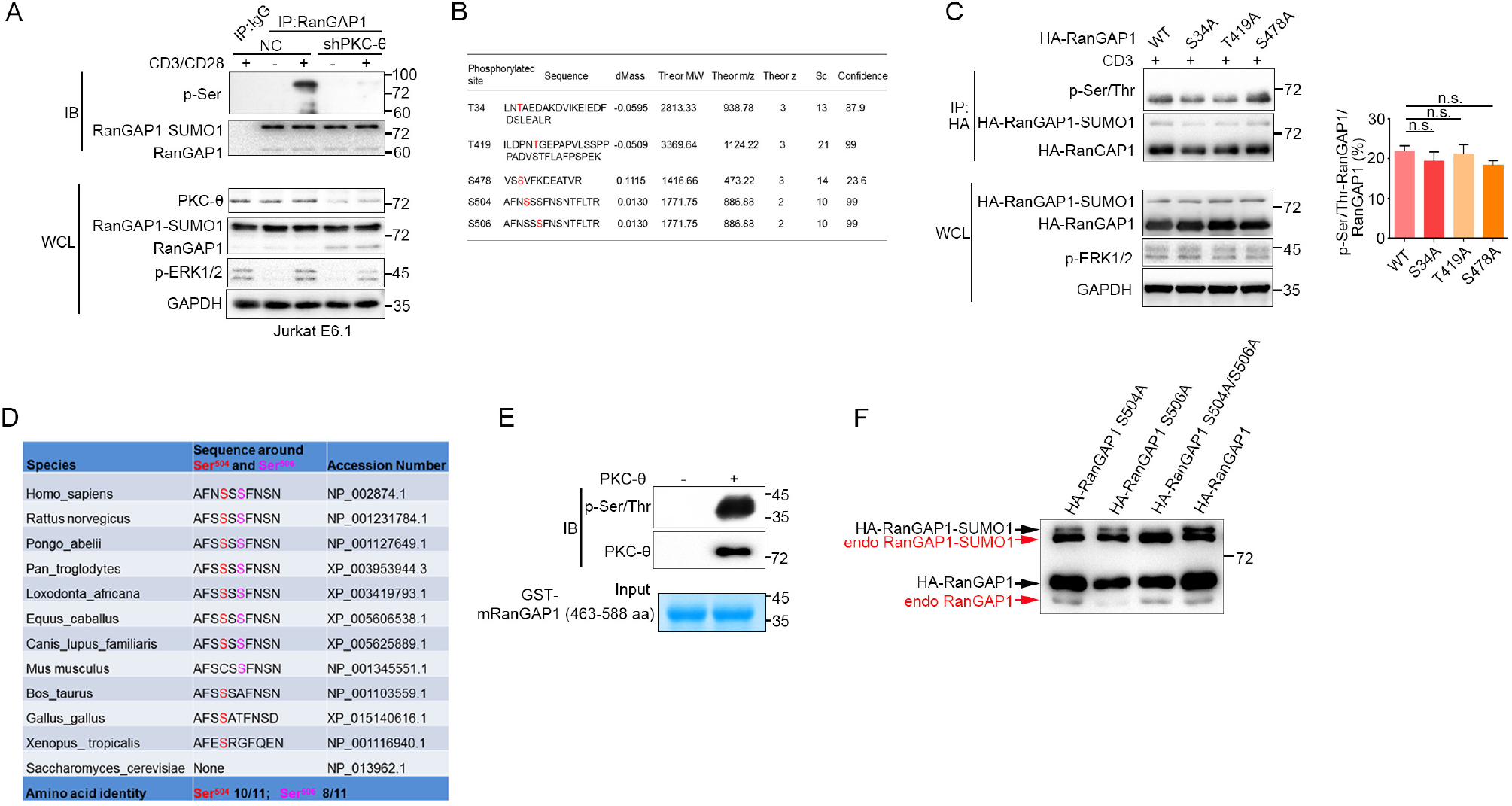
PKC-θ-dependent phosphorylation of RanGAP1. (**A**) Immunoblot analysis of the serine-phosphorylated RanGAP1 in stably transfected control (NC) or shPKC-θ-expressing Jurkat E6.1 T cells left unstimulated or stimulated with anti-CD3 plus anti-CD28. Asterisk indicates a nonspecific protein band. (**B**) Listing of 5 RanGAP1 Ser/Thr phosphorylation sites (red) identified by MS analysis of RanGAP1 phosphorylated *in vitro* by PKC-θ. (**C**) Immunoblot analysis of the phosphorylation of transfected wild-type or Ser-mutated HA-tagged RanGAP1 using a mixture of p-Ser- and p-Thr-specific antibodies in Jurkat-TAg cells, which were left unstimulated or stimulated for 15 min with anti-CD3 (left). Immunoblotting of the indicated proteins in WCL is shown at bottom. The ratio of phospho-RanGAP1 to immunoprecipitated RanGAP1 is shown in (right). Analysis is based on three biological replicates. n.s., not significant (one-way ANOVA with post hoc test). (**D**) Sequence alignment of RanGAP1 proteins from the indicated species, showing conservation of Ser^504^ and Ser^506^. (**E**) *In vitro* PKC-θ kinase assay using recombinant truncated GST-mouse mRanGAP1463-588 as a substrate. (**F**) Immunoblot analysis of both HA-RanGAP1 and endogenous RanGAP1 in WCL from Figure. *4F*, detected by RanGAP1 antibody. Data are representative of three (**A, C, E, F**) or two (**B**) biological replicates.

**Figure 5—figure supplement 1.**
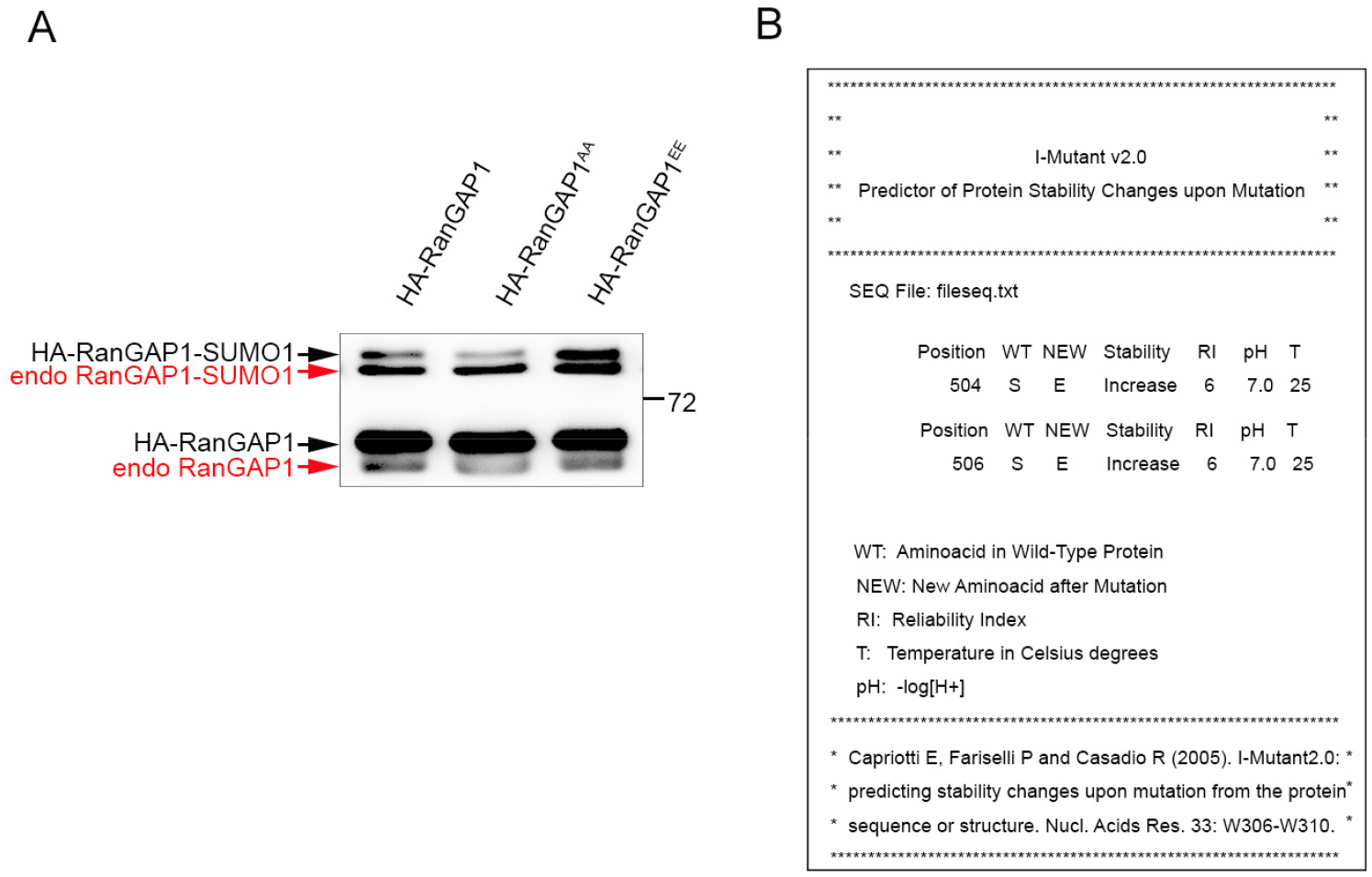
Characterization of the effect of Ser/Glu mutation on RanGAP1. (**A**) Immunoblot analysis of both HA-RanGAP1 and endogenous RanGAP1 in WCL from Figure 5(D), detected by RanGAP1 antibody. (**B**) Prediction of RanGAP1 stability changes upon mutating Ser^504^ and Ser^506^ to Glu. Predicted increase of protein stability due to mutations at serine 504 or serine 506 of RanGAP1 to glutamic acid as determined by i-Mutant2.0 (folding.biofold.org/i-mutant/i-mutant2.0.html). Data are representative of three (**A**) biological replicates.

**Figure 6—figure supplement 1.**
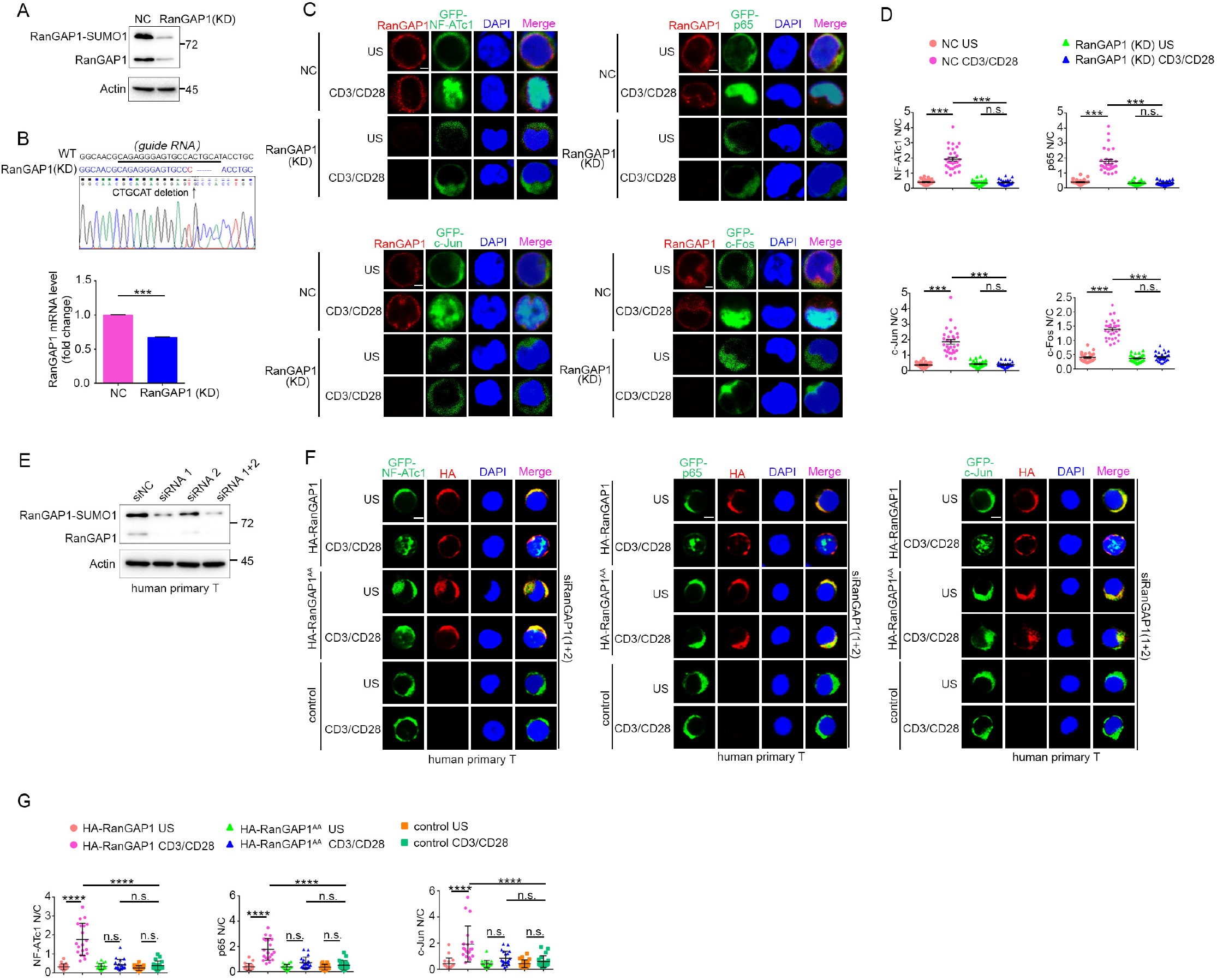
RanGAP1 is required for TCR-induced nuclear import of NF-ATc1, NF-κB, and AP-1. (**A**) RanGAP1 expression in negative control (NC, cells transfected with scrambled gRNA) Jurkat E6.1 T cells or cells with stable CRISPR/Cas9-edited RanGAP1 knockdown (KD). (**B**) Sequence alignment of RanGAP1 genomic fragment (WT) and the CRISPR/Cas9-edited RanGAP1 genomic fragment in the RanGAP1 (KD) Jurkat T cell line (top). The underlined sequence represents the guide RNA. The mRNA expression levels of RanGAP1 in NC and RanGAP1 (KD) T cells are shown at the bottom. (**C**) Confocal imaging showing the nuclear import of GFP-tagged NF-ATc1, p65 (NF-κB), c-Jun and c-Fos in NC and RanGAP1 (KD) T cells unstimulated or stimulated with anti-CD3 plus anti-CD28. Scale bars, 2 μm. These images are representative of ~30 cells analyzed in each group from two biological replicates. (**D**) Quantification of the nuclear import of GFP-tagged NF-ATc1, p65 (NF-κB), c-Jun and cFos as presented in (C). Each symbol represents an individual T cell. Horizontal lines indicate the mean ± s.e.m. n.s., not significant, \*\*\**P* < 0.001 (one-way ANOVA with post hoc test). (**E**) The knockdown effect of siRanGAP1 in human primary T cells. (**F**) Confocal imaging showing the nuclear import of transfected GFP-tagged NF-ATc1, p65 (NF-κB) or c-Jun in human primary T cells cotransfected with siNC or siRanGAP1 (siRNA 1+2), unstimulated or stimulated with anti-CD3 plus anti-CD28. Scale bars, 2 μm. These images are representative of ~30 cells analyzed in each group from two biological replicates. (**G**) Quantification of the nuclear import of GFP-tagged NF-ATc1, p65 (NF-κB), c-Jun as presented in (F). Each symbol represents an individual T cell. Horizontal lines indicate the mean ± s.e.m. n.s., not significant, *\*\*\*P* < 0.001, *\*\*\*\*p* < 0.0001 (one-way ANOVA with post hoc test). Data are representative of three (**A,E**) or two (**C, D, F, G**) biological replicates.

